# Human iPSC-derived brain pericytes exhibit differences in inflammatory activation compared to primary human brain pericytes

**DOI:** 10.1101/2024.09.16.613375

**Authors:** Samuel JC McCullough, Eliene Albers, Akshata Anchan, Jane Yu, Simon J O’Carroll, Bronwen Connor, E Scott Graham

## Abstract

**Background:** iPSC-derived cells are increasingly used to model complex diseases *in vitro* because they can be patient derived and can differentiate into any cell in the adult human body. Recent studies have demonstrated the generation of brain pericytes using a neural crest-based differentiation protocol. However, the inflammatory response of these iPSC-derived brain pericytes has not been investigated. We aimed to investigate the response of iPSC-derived brain pericytes to common inflammatory stimuli, thereby assessing the suitability of these cells to study inflammatory disease.

**Methods:** Brain pericytes were differentiated from iPSCs for 42 days. The expression of brain pericyte markers was assessed by RT-qPCR and immunofluorescent staining at days 0, 15, 21, and 42 of differentiation to validate the brain pericyte-like phenotype. Nuclear localisation of NFκB and STAT1 was assessed by immunofluorescence following IL-1β- and TNF-treatment in day 21 and day 42 iPSC-derived pericytes, and primary human pericytes. Cytometric bead array assessed the concentration of secreted inflammatory factors in the cell medium and phagocytosis was investigated using fluorescent carboxylated beads and flow cytometry.

**Results:** At day 42 of differentiation, but not at day 21, cells expressed brain pericyte markers. Generally, iPSC-derived pericytes lacked consistent responses to inflammatory treatment compared to primary human pericytes. Day 21 and 42 iPSC-derived pericytes exhibited a NFκB response to IL-1β treatment comparable to primary human pericytes. Day 21 iPSC-derived pericytes exhibited a STAT1 response with IL-1β treatment which was absent in day 42 cells, but present in a subset of primary human pericytes. TNF treatment presented similar NFκB responses between day 21 and 42 iPSC-derived and primary human pericytes, but a STAT1 response was again present in a subset of primary human pericytes which was absent in both day 21 and day 42 iPSC-derived pericytes. Numerous differences were observed in the secretion of cytokines and chemokines following treatment of iPSC-derived and primary human pericytes with IL-1β and TNF. iPSC-derived pericytes exhibited greater rates of phagocytosis than primary human pericytes.

**Conclusions:** With the increase in iPSC-derived cells in research, labs should undertake validation of lineage specificity when adapting an iPSC-derived differentiation protocol. In our hands, the inflammatory response of iPSC-derived pericytes was different to that of primary human pericytes, raising concern regarding the use of iPSC-derived pericytes to study neuroinflammatory disease.

**Graphical Abstract:** 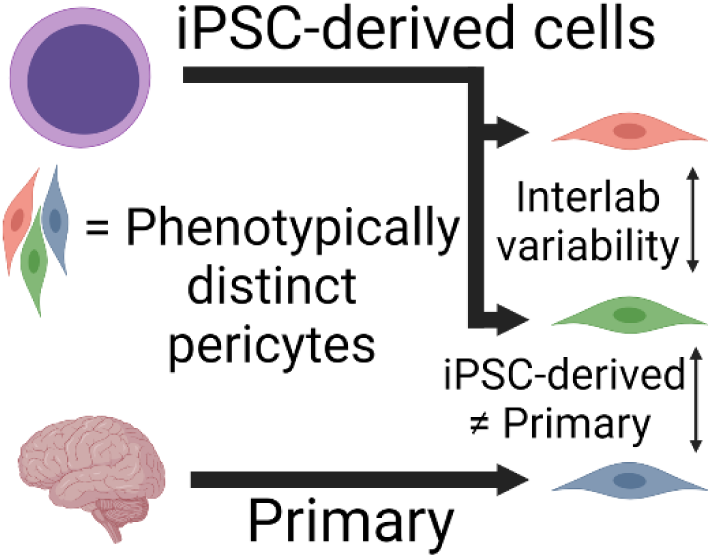

Brain pericytes can be generated from iPSCs. The work presented here shows the generation of phenotypically distinct pericytes from the original protocol, demonstrating the significant variability present within some iPSC differentiation protocols. Furthermore, functional differences are demonstrated between iPSC-derived brain pericytes and primary brain pericytes, revealing limitations in the use of iPSC-derived brain pericytes to model brain pericyte biology.

**Key Points:** *What is already known about this topic?:* Brain pericyte-like cells can be generated from induced pluripotent stem cells, however their responses to inflammatory stimuli has not been assessed.

*What does this study add?:* iPSC-derived brain pericytes exhibit different inflammatory responses compared to primary brain pericytes, showing that some iPSC-derived cell models are not appropriate for modelling all aspects of a cell’s biology. Furthermore, the iPSC-derived pericytes generated here were markedly different to those generated from the original article. It is therefore important for each lab to optimise the generation of iPSC-derived cell in their own hands to account for potential inter-lab variability.

## Introduction

Brain pericytes occupy a unique vascular niche within the basement membrane of brain endothelial cells. In the past, this physical niche has made adult brain pericytes difficult to study in isolation. Subsequently, the role of brain pericytes in blood-brain barrier (BBB) maintenance and the mediation of neuroinflammation has only recently been appreciated [1,2]. Adult brain pericytes are capable of responding to numerous inflammatory cytokines. Interleukin 1β (IL-1β) and tumour necrosis factor (TNF) activate downstream transcription factors to change pericyte function from vascular support to immunomodulatory [2]. Adult brain pericytes express the immune-modulating transcription factors nuclear factor kappa-light-chain-enhancer of activated B cells (NFκB) and signal transducer and activator of transcription 1 (STAT1), with the activation of each changing the brain pericyte’s inflammatory response while inducing the secretion of a plethora of inflammatory cytokines and chemokines [3]. Adult brain pericytes are also equipped to act as phagocytic cells when required, and their perivascular placement makes them an ideal second line of defence against potentially neurotoxic compounds [4,5].

Primary human brain pericytes are the gold standard for the *in vitro* study of brain pericyte function. However, primary human brain pericytes are difficult to obtain, typically requiring donation of post-mortem human tissue, resulting in a limited supply of these precious cells. To address this, human primary foetal (HPF) brain pericytes are commercially available, but it is unknown whether they recapitulate the phenotype of adult human brain pericytes. The most common platform to study brain pericytes is the use of animal models which can provide a powerful platform for pericyte research due to the added flexibility offered by genetically modified animals. However, several species-specific differences have been identified between mouse and human brain pericytes [2,6,7]. An example is the expression of *VTN* (the gene for vitronectin), which has previously been used as a marker for brain pericytes, which is now thought to be expressed in murine brain pericytes but not human pericytes [7]. Interestingly, vitronectin expression in mouse pericytes plays an important role in regulating the mouse BBB, demonstrating important species-specific differences between human and rodent brain pericytes [8]. Due to the limitations of current brain pericyte models, the generation of new *in vitro* models for studying brain pericyte biology is necessary.

The use of human induced pluripotent stem cell (iPSC)-derived pericytes is an exciting new platform for *in vitro* investigation of pericyte function. As iPSCs remain proliferative *in vitro*, the supply of iPSC-derived pericytes is greater than that obtained with adult primary human pericytes. Most importantly, iPSC-derived pericytes can be reprogrammed from patient-derived tissue. This allows for the creation of *in vitro* disease models to study changes in brain pericyte function under disease states, or to test the effect of pharmacological intervention on diseased pericytes. However, there are limitations in the use of iPSC-derived brain pericytes. When reprogrammed, the induced cells are rejuvenated at the transcriptional level, upregulating expression of foetal cell markers while aging signatures are erased [9,10]. As age is a strong risk factor for neurodegenerative diseases, iPSC-derived cells may lack properties acquired with age that contribute to the disease phenotype. Another limitation of iPSC-derived brain pericytes is the lack of a well-established brain-specific differentiation protocol. Pericyte-like cells have been generated from iPSCs since 2009, however, most of these protocols used a mesodermal differentiation protocol, generating non-brain pericytes (Table S1). Brain pericytes are derived from neural crest stem cells (NCSCs) and display transcriptional differences to non-brain pericytes [1,11]. Recently, brain pericyte differentiation protocols have been published which exhibit characteristic brain pericyte marker expression, as well as some functional aspects of brain pericyte biology such as supporting endothelial barrier function ^[12-^

^15]^. However, these studies have yet to study the inflammatory responses of iPSC-derived brain pericytes. Given that some differentiation protocols require weeks of culture before generating mature pericytes, it is important that the end-product is fit for purpose.

This study aimed to determine the suitability of iPSC-derived brain pericytes for studying inflammatory disease. A recently published brain pericyte differentiation protocol was used to generate brain pericyte-like cells [14]. We hypothesised that iPSC-derived brain pericytes would produce comparable responses to that of HPF pericytes, thereby justifying the use of iPSC-derived brain pericytes as an alternative platform to study brain pericyte involvement in the inflammatory response.

## Materials and methods

### Induced pluripotent stem cell culture

The IPS(IMR-90)-4 (WiCell) iPSC line (known here as IMR-90) was used to generate iPSC-derived pericytes. The IMR-90 iPSC line was reprogrammed with a lentiviral vector containing OCT3/4, SOX2, NANOG, and LIN28. IMR-90 iPSCs were routinely cultured in extracellular matrix (ECM) gel-coated 6-well plates (8.75µg/cm^2^; catalogue#: E6909, Sigma-Aldrich). ECM gel was diluted to 8.75µg/cm^2^ in 1mL/well DMEM/F12 and spread evenly across the bottom of the 6-well plate. The ECM gel was left at 37°C for at least 1 hour before use. The iPSC lines were maintained with daily refreshment of Essential 8 medium (catalogue#: A1517001, Thermo Fisher Scientific). To passage iPSC lines, spent medium was removed and the cells were washed with 1mL of warmed phosphate buffered saline (PBS). To detach the cells from the plate, 1mL of Versene (catalogue#: 15040066, Thermo Fisher Scientific) was added and the cells were placed at 37°C for 3 minutes. The cells were then viewed under a bright field microscope to confirm the loosening of colony attachment through feathered colony borders. If cells were yet to detach, they were incubated at 37°C for additional 1-minute increments until detachment was seen, or until the cells had been incubated for a total of 5 minutes. The Versene was carefully removed and 1mL of fresh warmed iPSC medium was added to each well to wash off the colonies. When passaging the iPSCs, the cell suspension was triturated a maximum of three times to keep iPSC colonies together. This 1mL of medium was added to 11mL of fresh warmed iPSC medium (this volume passages a single well 1:6 and was scaled up if required). The iPSCs were then seeded into six wells of an ECM-coated 6-well plate with a final volume of 2mL for a 1:6 split. If the iPSCs were to be frozen, the same steps as above were followed until removal of Versene, where the cells were instead resuspended in 1mL mFresR (catalogue#: 05855, Stemcell Technologies) and placed in a cryovial for cryostorage.

When plating iPSCs for experimentation, Accutase (catalogue#: 07922, Stemcell Technologies) was used as a dissociation reagent rather than Versene. This promoted a single cells suspension, as opposed to the preservation of iPSC colonies. Accutase was incubated with the iPSCs for 3 minutes, or until cell rounding could be seen under a brightfield microscope. The Accutase was triturated three times to detach cells and the cells were collected in spent medium to dilute the Accutase at least 1:2. This cell suspension was centrifuged at 300xg for 4 minutes, the supernatant was discarded, and the iPSCs were suspended in Essential 8 medium supplemented with 10μM Y27632. Y27632 is a ROCK inhibitor which prevented apoptosis in the dissociated iPSCs. The iPSCs were then plated out at the desired density for the experimental application.

### Primary pericyte cell culture

Two different primary pericyte cell suppliers were used in this study: Creative Bioarray Primary Human Brain Cortex Pericyte Cells (catalogue#: CSC-C4387X, Creative Bioarray), and ScienCell Human Brain Vascular Pericytes (catalogue#: 1200, Sciencell). The Creative Bioarray Primary Human Brain Cortex Pericyte Cells were used as a positive control for characterisation of the iPSC-derived brain pericytes, including RT-qPCR and immunocytochemistry related to characterisation. ScienCell Human Brain Vascular Pericytes were used for immunocytochemistry related to inflammatory treatments, cytometric bead array, and phagocytosis experiments in this study. Given the foetal origins of both cell lines, they are referred to collectively as “HPF pericytes”. Cell culture methods were similar between the two HPF pericyte lines, though ScienCell primary pericyte cell culture flasks were coated with poly-l-lysine (2µg/cm^2^; catalogue#: P1274, Sigma-Aldrich), and different medium was used for each culture. Poly-l-lysine was diluted to 2µg/cm^2^ in sterile milliQ water and added to the culture vessel. Coated flasks/plates were incubated at 37°C for at least 1 hour before use. Creative Bioarray primary pericytes were cultured in DMEM/F12 (catalogue#: 11330032, Thermo Fisher Scientific) with 10% FBS (catalogue#: FBSF, Moregate Biotech) and 1% Penicillin-Streptomycin (catalogue#: 15070063, Thermo Fisher Scientific). ScienCell HPF pericytes were cultured in ScienCell complete pericyte medium (catalogue#: 1201, Sciencell).

HPF pericytes were typically cultured in T75 flasks, where cell culture medium was routinely refreshed every 2-3 days depending on cell density. Cells were passaged at 90% confluency. To passage HPF pericytes, the cells were washed with warm PBS before 0.025% EDTA trypsin was added to the flask. HPF pericytes were placed in a 37°C incubator for 2 minutes with trypsin to promote cell detachment. Detachment was observed as cell rounding under a light microscope. With confirmation of cell detachment, HPF pericytes were washed off the flask with spent medium which diluted the trypsin at least 1:4. The HPF pericyte cell suspension was spun down at 300g for 5 minutes, the supernatant was removed, and cells were resuspended and counted before seeding / cell freezing. For routine passaging, HPF pericytes were seeded in T75 flasks at 5,000 cells/cm^2^. For experiments, HPF pericytes were seeded in 96-well plates at 15,000 cells/cm^2^, or 24 well plates at 5000 cells/cm^2^. HPF pericytes could also be frozen at 1,000,000 cells/vial. Cells were frozen in freezing medium (50% culture medium, 40% FBS, 10% DMSO) and stored at -80°C before being transferred to liquid nitrogen for long-term storage.

### Differentiating iPSCs into brain pericyte-like cells

A schematic overview of the differentiation protocol is provided in Figure 1A. The protocol to differentiate iPSCs into brain pericyte-like cells was first published by Stebbins et al., 2019. This protocol primes iPSCs down a neural crest stem cell (NCSC) lineage, before isolating NCSCs and differentiating them into brain pericyte-like cells. The priming of iPSCs towards a neural crest stem cell lineage was initiated 24 hours after dissociation and replating of iPSCs at 875,000 cells/well in a 6-well plate. The iPSCs were dissociated with Accutase to generate a single cell suspension and were plated in Essential 8 medium with Y27632 as described in the cell culture methods. After 24 hours, the medium was changed to neural crest stem cell medium (known as E6-CSFD; see Table S2). Differentiating cells are maintained in E6-CSFD with medium refreshed daily and cells passaged 1:6 at 90-100% confluency. This continued until day 15, where NCSCs were positively isolated from the mixed culture using p75-conjugated magnetic beads (catalogue#: 130-097-127, Miltenyi Biotec). Day 15 cells were collected with Accutase, using general iPSC culture methods. To perform the isolation, the mixed NCSC culture of one well of a 6-well plate was suspended and the cells were counted to provide an estimate of the number of cells in each well. An appropriate number of wells were suspended and passed through a 40µm cell strainer such that less than 20,000,000 cells were run through the isolation at once (catalogue#: 22-363-547, Thermo Fisher Scientific) to prevent clumps in the suspension before being spun down at 300g for 4 minutes. Cells were suspended in 80µL / 10,000,000 cells FACS buffer (PBS, 0.5% BSA, 2nM EDTA), and 20µL / 10,000,000 cells p75-conjugated magnetic bead was added. This bead cocktail was incubated at 4°C for 15 minutes, flicking the tube halfway through the incubation to prevent cells from adhering to the tube. Following this incubation, 2mL FACS buffer was added, and the cells were spun down at 4°C, 300g for 4 minutes. During this time, the MACS column (catalogue#: 130-042-401, Miltenyi Biotec) was placed in a MidiMACS separator (catalogue#: 130-042-302, Miltenyi Biotec) and primed by adding 500µL FACS buffer. The labelled cells were resuspended in 500µL FACS buffer and placed in the MACS column. The column was washed with 1mL FACS buffer three times, collecting p75-negative cells if appropriate for the experiment, before being removed from the MidiMACS separator and removing the p75-positive cells in the column. These cells are spun down, resuspended in 500µL FACS buffer, and passed through another column using the same methods as above. Following this second isolation, p75-positive cells are resuspended in pericyte differentiation medium (known as E6+10% FBS; see Table S3), counted, and seeded into uncoated 6-well plates at a density of 500,000 cells/well (or ∼50,000 cells/cm^2^). The differentiating iPSC-derived pericytes were maintained in E6+10% FBS with medium refreshed daily and cells passaged 1:2 at 90-100% confluency until day 43. Pericyte-like cell senescence was observed soon after D43.

**Figure 1:**
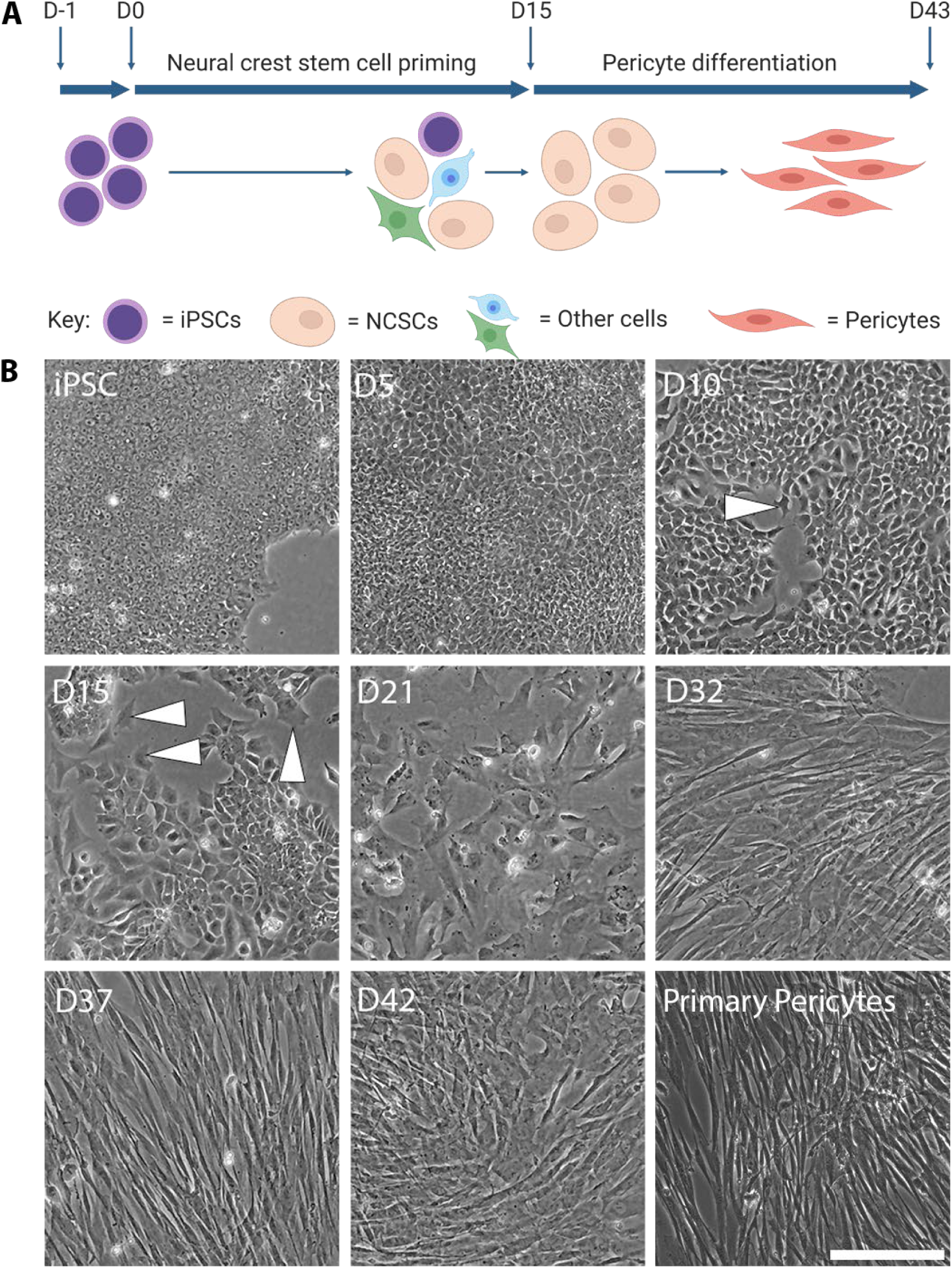
iPSCs are differentiated into brain pericyte-like cells. (A) Schematic of the brain pericyte differentiation protocol developed by Stebbins et al [14]. (B) Images of differentiating cells from iPSC to day 42 (D42) of pericyte differentiation. NCSC priming (D0-D15) results in a heterogeneous population of cells including larger cells at the colony border (white arrows). NCSCs are isolated and grown in pericyte differentiation medium, at which point a more homogenous population of cells can be seen (D21). Differentiating cells acquire an elongated morphology over the period of pericyte differentiation (D15-D42). This morphology is comparable to the morphology seen in human primary foetal pericytes. Scale bar = 200μm.

### Investigating transcriptional expression with RT-qPCR

RNA was extracted from cells using the Macherey-Nagel Nucleospin RNA purification kit (catalogue#: 740955.50, Macherey-Nagel) following the manufacturers methods. A DNA copy (cDNA) was synthesised from the extracted RNA such that it becomes compatible for RT-qPCR. Standard methods using the SuperScript IV First-Strand Synthesis System (catalogue#: 18091050, Thermo Fisher Scientific) were followed for this procedure. To perform RT-qPCR, 5μL of TaqMan Gene Expression Master Mix (Catalogue#: 4369016, Thermo Fisher Scientific) was added to an RNase-free tube and was mixed with 0.5μL/sample of 18S-VIC primer (catalogue#: 4448486, assay ID: Hs03003631_g1, Thermo Fisher Scientific) and 0.5μL/sample of target gene primer to make the reaction mix (see Table S4 for details of primers used in this study). 6μL of reaction mix and 4μL cDNA (equivalent of 4µg) was added to a 384-well qPCR plate for each sample in triplicate. Duplex qPCR reactions were run on a Quantstudio 12 using 18 S rRNA as the internal control to generate amplification curves. 18 S rRNA was chosen as the internal control due to its consistent expression across cell types, including reprogrammed cells. Amplification curves from each sample were analysed using Applied Biosystems QuantStudio 5 Real-Time PCR System to generate ΔCt values.

### Investigating protein expression with immunocytochemistry

Cells were seeded at 15,000 cells/cm^2^ in 96-well plates for inflammatory transcription factor immunocytochemistry, or at 5000 cells/cm^2^ on 13mm-coverslips in 24 well plates. Cell culture samples were treated with inflammatory factors or vehicle (see Table S5) for 1 hour before fixation with 4% paraformaldehyde for 10-15 minutes. Inflammatory concentrations and treatment times are based on previous work describing robust stimulation in primary human brain pericytes [3]. Cells were washed with PBS and permeabilised with PBS-triton (0.1%) to reveal intracellular epitopes. Primary antibodies were diluted in immunobuffer (PBS + 1% foetal donkey serum) and were incubated with the cells at 4°C overnight (see Table S6 for details of antibody dilutions used in this study). The following day, the cells were washed 2-3 times with PBS before secondary antibodies diluted in immunobuffer with Hoechst 33342 were added to the cells and incubated overnight at 4°C. The following day, the stained cells were washed 2-3 times with PBS. For inflammatory transcription factor immunocytochemistry, stained cells remained in PBS in a 96-well plate to be imaged. Cells that were grown on 13mm-coverslips in 24-well plates were carefully removed and placed on Permafrost microscope slides (catalogue#: M642, Cardinal Health 200) with ProLong Diamond Antifade Mountant with DAPI (catalogue#: P36966, Thermo Fisher Scientific). Slides were left covered at room temperature overnight to allow the mountant to cure before imaging.

### Cytometric bead array to study secreted factors

Cytometric bead array (CBA) experiments investigated changes in iPSC-derived pericyte and HPF pericyte secretion in response to treatments. Experiments were performed as described previously [16]. A list of CBA kits used in this study can be found in Table S7.

### Using fluorescent carboxylated beads to study pericyte phagocytosis

50µL of Fluoresbrite YG carboxylated microspheres (1μm, catalogue#: 15702-10, Polysciences) diluted in pericyte medium was spiked into pericyte cell medium in 24-well plates. The beads were diluted to generate a final effector-target ratio (bead to cell ratio) of 1300:1. To study phagocytosis through fluorescent imaging, cells were fixed and permeabilised as described in the immunocytochemistry methods. Cells were then incubated with Hoechst 33342 (1:10,000) at 4°C overnight. The following day, the cells were washed three times with PBS and were imaged on the EVOS FL Auto.

To study brain pericyte phagocytosis through flow cytometry, the carboxylated microspheres were added to the cell culture as above. Cells were (pre-)treated with inflammatory treatments as described in the relevant methods section. After incubation with carboxylated microsphere for 24 hours, the cells were harvested with trypsin. Trypsin cleaves cell surface proteins, removing surface-bound beads. Cells were centrifuged at 300xg and were resuspended in FACS buffer as a wash step (PBS, 0.5% BSA, 2nM EDTA). Brain pericytes were only washed once to conserve cell viability. The cell suspension was pelleted again, and the supernatant was removed before the cells were resuspended in fresh FACS buffer. Cells were stored on ice until being run on the Accuri C6 flow cytometer. Brain pericytes without bead treatment were run to show the auto-fluorescence of each pericyte population alone. This auto-fluorescent threshold was used to determine whether fluorescent bead was phagocytosed by pericytes, revealing the percentage of phagocytic cells present in the population. The mean fluorescent intensity (MFI) of phagocytic cells was also assessed. The gating strategy used to generate histo-plots can be found in the supplementary material (Figure S1)

### Imaging methods to capture and quantify light and fluorescent images

Phase and brightfield images were taken of live cells and were captured on either the EVOS XL CORE (catalogue#: AMEX1000, Thermo Fisher Scientific) or the EVOS FL (catalogue#: 12-563-460, Thermo Fisher Scientific) using a 10x objective. Fluorescent images were captured on the EVOS FL Auto (catalogue#: AMF7000, Thermo Fisher Scientific) to characterise the iPSC-derived pericytes, image phagocytosis assays, or to determine inflammatory transcription factor immunocytochemistry specifically in the iPSC-derived cells. Samples were imaged at 10x and 20x. Inflammatory transcription factor immunocytochemistry in HPF pericytes was imaged at 10x on the Molecular Devices ImageXpress Micro XLS for high-throughput screening. To quantify inflammatory transcription factor translocation in iPSC-derived pericytes, an ImageJ macro was developed to measure the fluorescence that co-localised with each nucleus (see Table S8). This was thresholded to identify each cell as either positive or negative for nuclear signal. To assess inflammatory transcription factor translocation in HPF pericytes, images taken on the ImageXpress Micro XLS were analysed using the “Multi Wavelength Translocation” software package in MetaXpress to identify each cell as either positive or negative for nuclear signal.

### Statistical analysis

Significant changes between groups for qPCR experiments was not assessed as this did not address our research question: whether the genes were expressed or not. For concentration-response data, significant change from vehicle-treated samples was determined using a one-way ANOVA with Bonferroni’s multiple comparisons test. In some cases, statistical significance was not reached due to a lack of experimental repeats or variation in the response – often seen in the iPSC-derived pericytes.

## Results

### iPSCs can be differentiated into brain pericyte-like cells

As brain pericytes arise from cells of the neural crest, iPSCs are first primed down a NCSC lineage (Figure 1A). Phase images show that iPSCs are small cells (∼12µm diameter) with limited cytoplasm. These cells grow in colonies with a defined border (Figure 1B, iPSC). During NCSC priming, cells lose their iPSC morphology and become larger, migratory cells (Figure 1B, white arrows). By the end of NCSC priming on day 15, a morphologically heterogeneous population of cells exists; some compact and iPSC-like, others larger and migratory (Figure 1B, white arrows). To isolate the NCSC population, the cells were processed using MACS technology, utilising the NCSC marker p75 conjugated to magnetic beads. This process generated a 99.6% pure population of NCSCs as measured by immunocytochemistry for the NCSC marker p75 (see Figure S2). Additionally, the expression of NCSC markers *PAX3* and *PAX7* further confirmed a NCSC phenotype (Figure S3). The p75-positive NCSCs were plated in pericyte differentiation medium (E6+10%FBS), marking the initiation of pericyte differentiation. On day 21, a homogenous population of polygonal cells was observed by phase contrast imaging (Figure 1B, D21). From day 21-day 37 of pericyte differentiation, the cell morphology changed from polygonal to elongated (Figure 1B, D21-D37). This elongated morphology was maintained for the remaining duration of pericyte differentiation and was comparable to the morphology of HPF pericytes (Figure 1B, D42; primary pericytes). By day 42, differentiating cells can be seen growing on top of each other. This suggests that these cells have lost the contact inhibition expected from adult brain pericytes *in vitro*. The cells stopped proliferating between D42-D50, suggesting a similar *in vitro* response to primary pericytes, which also senesce with time [17].

Expression of several pluripotency markers were investigated over the course of differentiation to ensure loss of pluripotency with cell differentiation. *OCT3/4*, *SOX2* and *NANOG* were highly expressed in iPSCs with average ΔCt values of 12.5, 16.7, and 13.9 respectively (Figure 2A). By day 42 of pericyte differentiation, these ΔCt values had risen to 23.5, 21.4, and 16.3 respectively. While these higher ΔCt values indicate lower levels of gene expression, it also shows that gene expression is still present in the iPSC-derived brain pericytes. Interestingly however, expression of *OCT3/4*, *SOX2* and *NANOG* was also present in HPF pericytes with average ΔCt values of 22.7, 26.4, and 27.8 (Figure 2A). Protein expression of OCT3/4, SOX2, and NANOG was exhibited in iPSCs, but was not detected in iPSC-derived pericytes throughout differentiation, suggesting a mechanism to inhibit protein expression of these pluripotency transcription factors in these cells (Figure 2B).

**Figure 2:**
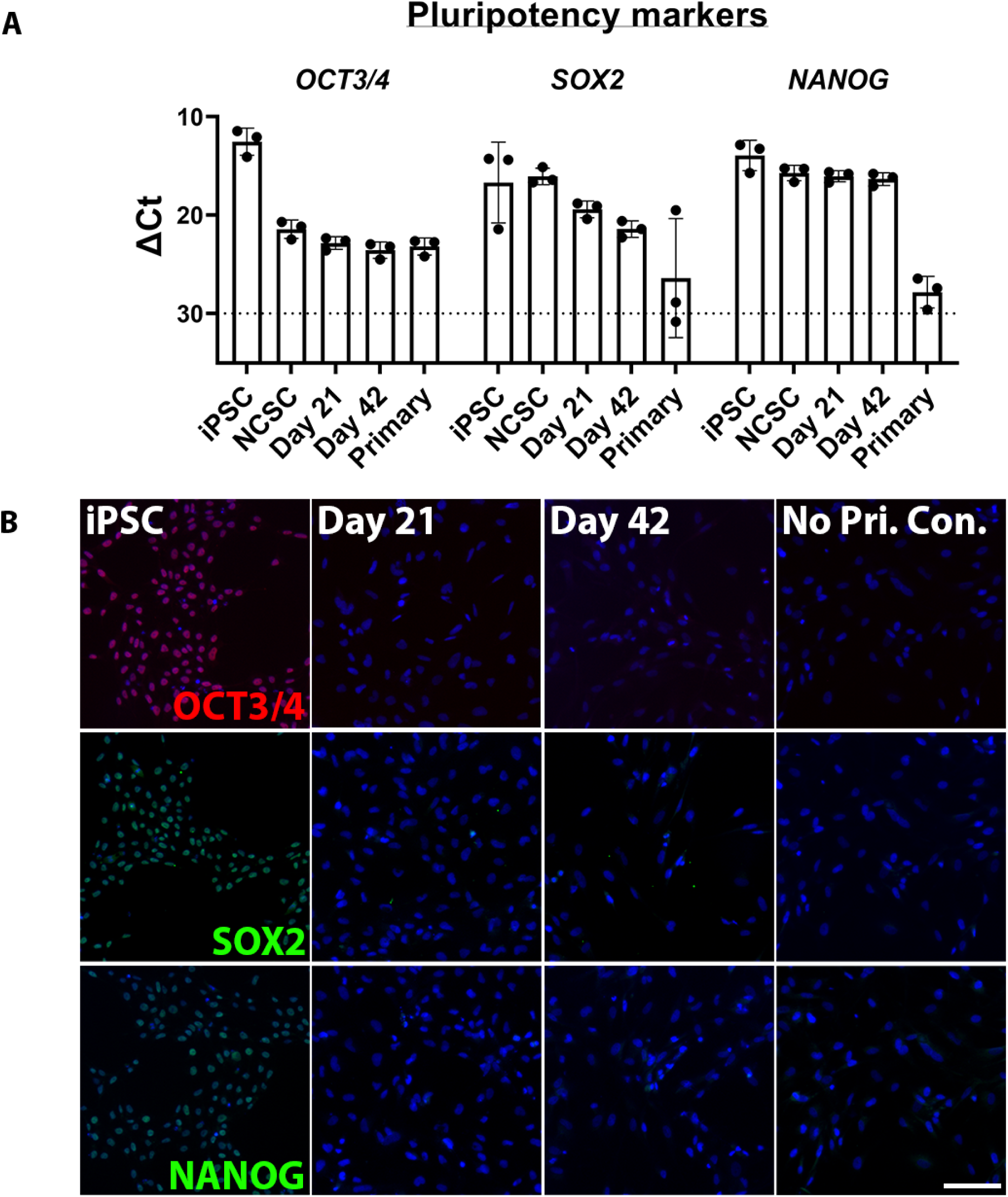
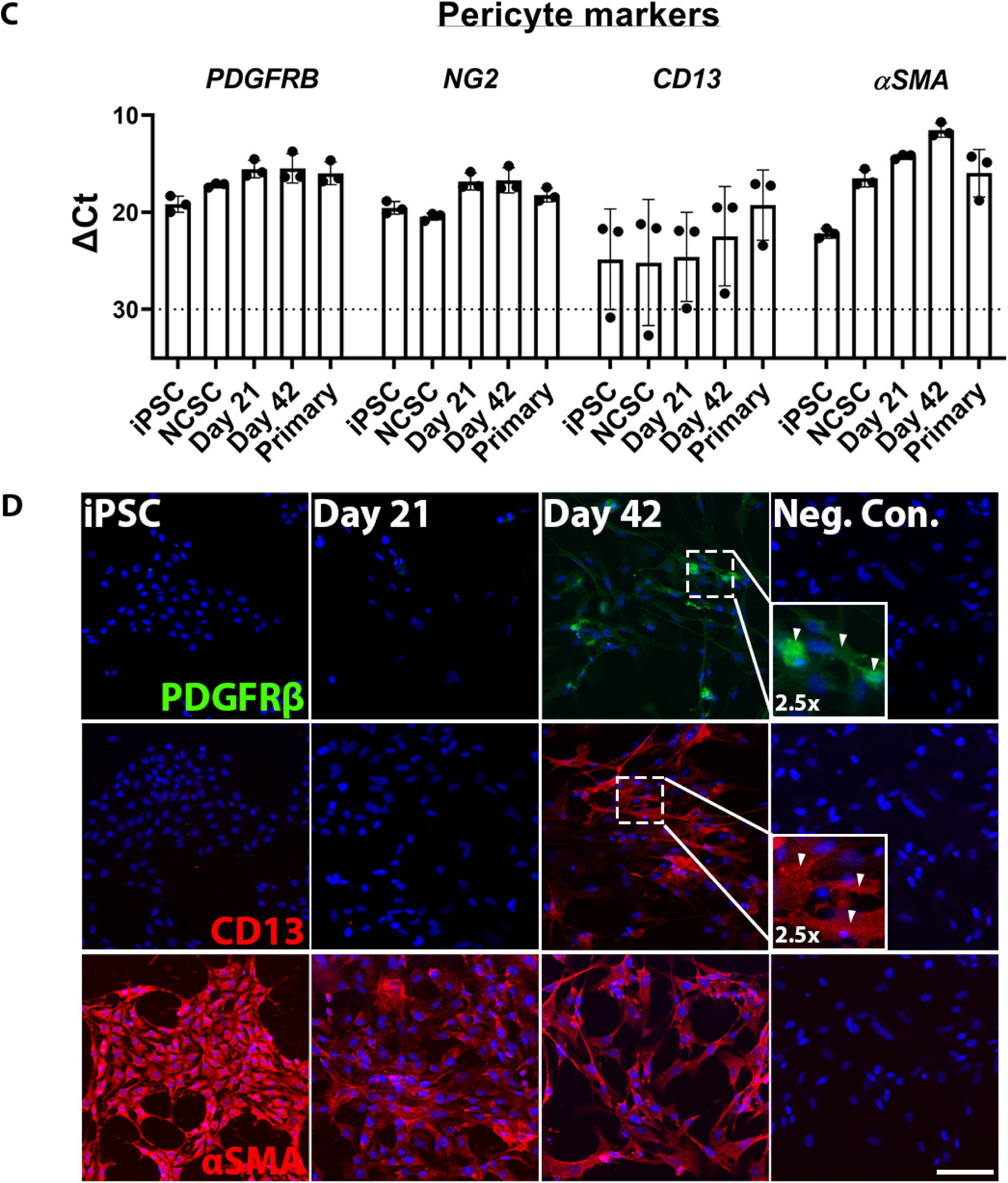
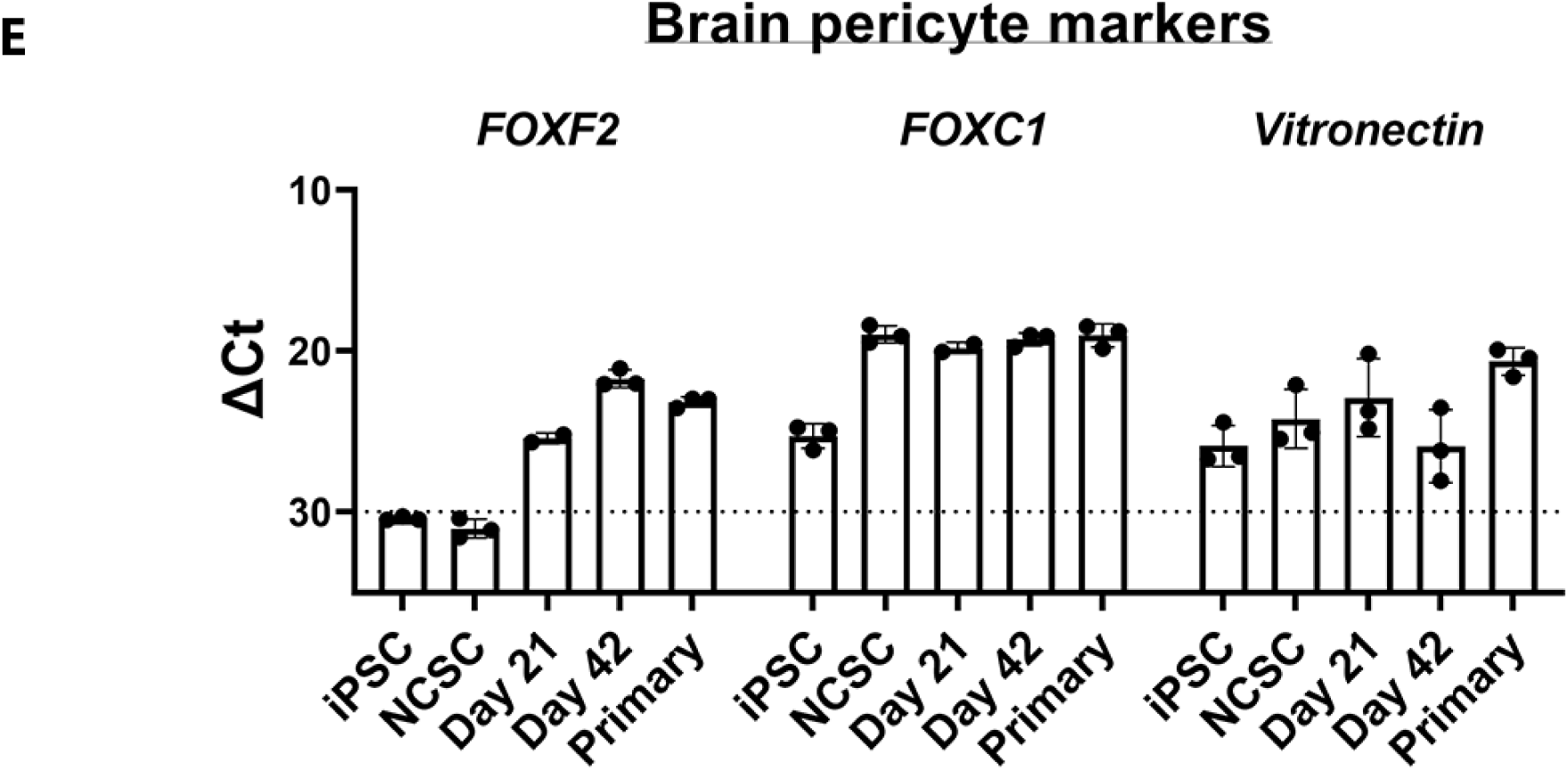
iPSC-derived brain pericytes exhibit primary human brain pericyte-like characteristics. (A) RT-qPCR shows high levels of expression of pluripotency gene expression (OCT3/4, SOX2, and NANOG) in iPSCs, with lower levels of expression seen in iPSC-derived pericytes. These pluripotency genes are also expressed at low levels in human primary foetal (HPF) pericytes. (B) Immunofluorescence images demonstrating pluripotency marker protein expression (OCT3/4, SOX2, and NANOG) in iPSCs, but not iPSC-derived pericytes. (C) RT-qPCR shows gene expression of pericyte markers (PDGFRβ, NG2, CD13, and αSMA) in iPSC-derived and HPF pericytes. (D) Immunofluorescence images demonstrating pericyte marker protein expression (PDGFRβ, CD13, and αSMA) in day 42 iPSC-derived pericytes. High levels of αSMA protein expression are observed in iPSCs and day 21 iPSC-derived pericytes (E) RT-qPCR shows gene expression of brain-specific pericyte markers (FOXF2, FOXC1, and vitronectin) in day 42 iPSC-derived pericytes and HPF pericytes. Gene expression of FOXF2 is absent in iPSCs and NCSCs. The dotted line on all RT-qPCR graphs indicates a ΔCt of 30, demonstrating the minimum ΔCt threshold of expression in these experiments. “Neg. Con.” refers to the negative control that didn’t receive primary antibody. Scale bar = 100μm in all images. Error bars represent standard deviation between the three experimental repeats.

Expression of the classical pericyte markers platelet-derived growth factor receptor-β (PDGFRβ), neural-glial antigen-2 (NG2), CD13, and α smooth muscle actin (αSMA) were investigated by RT-qPCR and immunocytochemistry to assess the generation of a pericyte-like phenotype (Figure 2C/D). Gene expression of these markers was detected in iPSCs, which exhibited average ΔCt values of 19.1, 19.5, 24.8, and 22.2 respectively (Figure 2C). Expression of these markers increased across pericyte differentiation, peaking at day 42 where the average ΔCt values were 15.5, 16.7, 22.4, and 11.5 respectively. These values are comparable to the level of expression seen in HPF pericytes, which demonstrated average ΔCt values of 15.6, 18.2, 19.3, and 16.0 for these genes respectively (Figure 2C). PDGFRβ immunocytochemistry showed PDGFRβ protein expression in day 42 iPSC-derived pericytes, but not iPSCs or day 21 iPSC-derived pericytes (Figure 2D). This is a considerable difference to the findings from the original protocol where the differentiating cells were considered pericyte-like by day 21 [14]. Negligible CD13 staining was seen prior to day 42, at which point CD13 can be seen in many but not all iPSC-derived pericytes in culture (Figure 2D). αSMA was found to be expressed by iPSCs, as well as all differentiating pericyte-like cells (Figure 2D). With the high levels of αSMA, we wanted to confirm that the generated cells were not smooth muscle cells by investigating expression of the pericyte/fibroblast marker *ABCC9* (also known as SUR2). *ABCC9* was present in iPSC-derived pericyte cultures, indicated that these are not smooth muscle cells (see Figure S4).

Having investigated the expression of classical pericyte markers, we assessed whether the protocol generated brain specific pericyte-like cells. To achieve this, the expression of brain pericyte markers *FOXF2*, *FOXC1*, and *VTN* (vitronectin) was assessed by RT-qPCR (Figure 2E). Though *VTN* was suggested to be a mouse pericyte marker in a recent single-cell RNA Seq study, given its importance in mouse brain pericyte biology, we wanted to confirm its presence or absence in human brain pericytes [7,8]. Expression of *FOXF2* was not detected in iPSCs or NCSCs (ΔCt of >30). However, expression was seen over the course of pericyte differentiation with day 42 iPSC-derived pericytes exhibiting a ΔCt of 21.7 (Figure 2E). This is comparable to the expression seen in HPF pericytes which exhibit a ΔCt of 23.2. These data demonstrate the upregulation of *FOXF2* over the course of pericyte differentiation. *FOXC1* expression was demonstrated at low levels in iPSCs with a ΔCt of 25.3 which increased to high expression in NCSCs with a ΔCt of 19.0 (Figure 2E). This high level of expression was maintained over the course of pericyte differentiation and is again comparable to expression demonstrated by HPF pericytes which exhibited a ΔCt of 19.0. *VTN* was found to be expressed in iPSCs with a ΔCt value of 25.9, increasing over differentiation until day 21 where the ΔCt value was 22.9 (Figure 2E). From day 21 to day 42 of pericyte differentiation, *VTN* expression decreases to the level of expression seen in iPSCs with a ΔCt value of 25.9. HPF pericytes demonstrated moderate expression of *VTN* with a ΔCt of 21.0, more like the level of expression seen in day 21 iPSC-derived pericytes. Together, these data show that day 42 iPSC-derived pericytes express brain pericyte cell markers, indicating that a brain pericyte-like phenotype may be induced in these cells.

### iPSC-derived brain pericytes exhibit different STAT1 responses to IL-1β and TNF treatment than human primary foetal brain pericytes

The pro-inflammatory cytokine IL-1β is known to cause NFκB translocation to the nucleus in primary adult human brain pericytes [3]. Both iPSC-derived and HPF pericytes demonstrated nuclear translocation of NFκB at comparable concentrations of IL-1β. The mean EC50 values were 7.57pM, 7.88pM, and 1.75pM in iPSC-derived day 21, iPSC-derived day 42, and HPF pericytes respectively (Figure 3A-F). This data indicates that the iPSC-derived pericytes possess the NFκB intracellular signalling cascade. However, the standard deviation between technical replicates makes apparent the heterogeneity within the iPSC-derived populations. The iPSC-derived pericytes also exhibited a smaller proportion of responsive cells. An average of 56.8% of day 21 iPSC-derived pericytes showed NFκB translocation, and 58.1% by day 42 (Figure 3A-F). A higher proportion of HPF pericytes responded, with an average of 82.6% of cells demonstrating NFκB translocation (Figure 3A-F). While this data suggests that both iPSC-derived and HPF pericytes possess the ability to signal through the IL-1β∷NFκB signalling cascade, HPF pericytes exhibit greater sensitivity and consistency than iPSC-derived pericytes.

**Figure 3:**
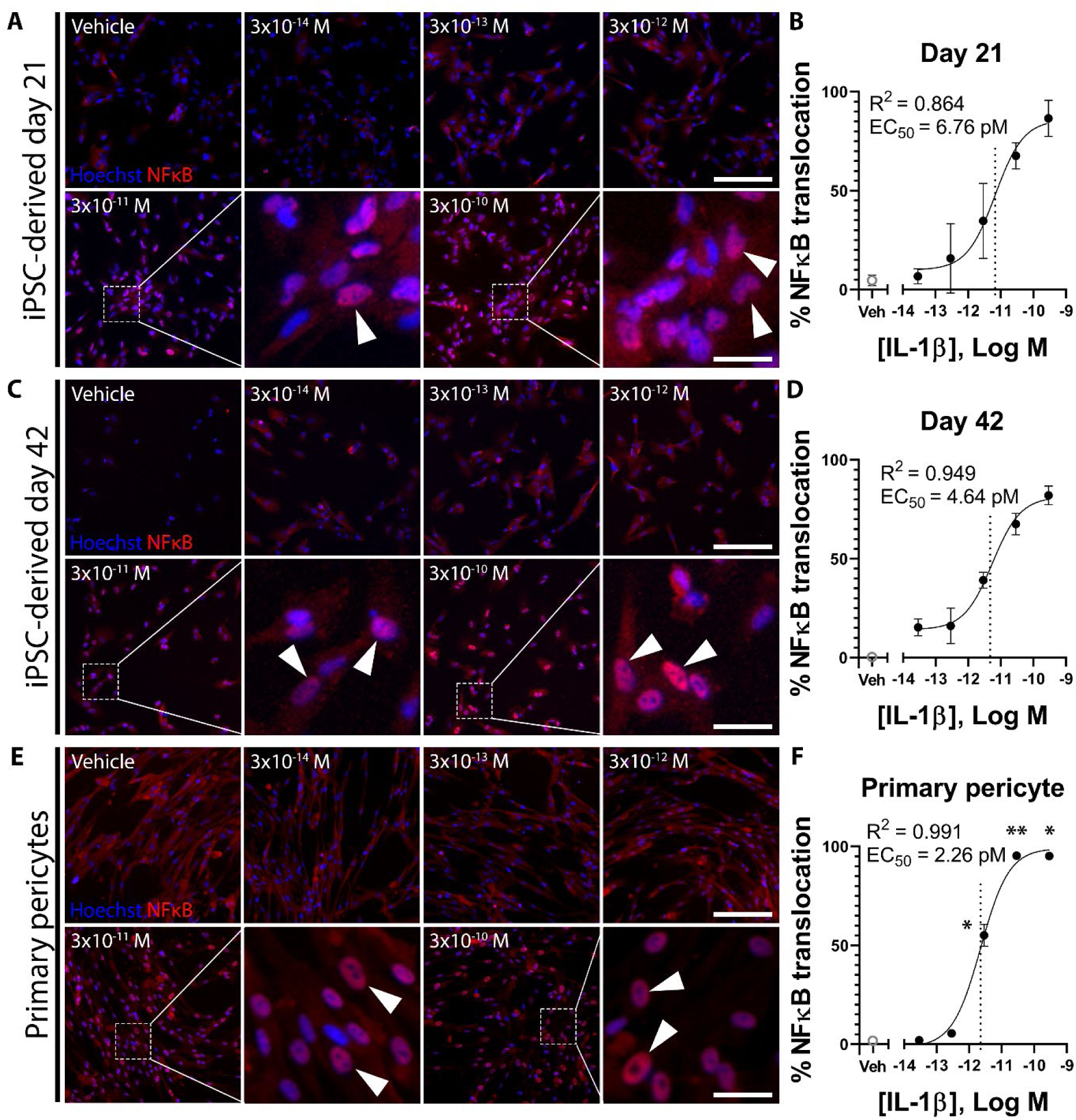
NFκB translocates to the nucleus with IL-1β treatment in day 21 and day 42 iPSC-derived pericytes and primary human pericytes. Immunofluorescence images demonstrating nuclear translocation of NFκB in response to increasing concentrations of IL-1β in day 21 iPSC-derived pericytes (A), day 42 iPSC-derived pericytes (C), and human primary foetal (HPF) brain pericytes (E). Images of iPSC-derived pericytes are quantified using an ImageJ macro to generate concentration-response curves (B, D) which demonstrate NFκB translocation at an EC_50_ of 6.76pM in day 21 iPSC-derived pericytes (B, dotted line), and 4.64pM in day 42 iPSC-derived pericytes (dotted line, D). Images of HPF pericytes are quantified using MetaXpress, showing an EC_50_ of 2.26pM (dotted line, F). Data presented is one representative experiment of two experimental repeats with the iPSC-derived pericytes, and one representative experiment of four experimental repeats with the HPF pericytes (see Table S9). Scale bar = 200µm with exception of 40µm for all further magnified images. White arrows indicate nuclear NFκB. Error bars represent standard deviation. Statistical significance was determined using one-way ANOVA with Bonferroni’s multiple comparisons test. * = P<0.05, ** = P<0.01, *** = P<0.001.

In addition to investigating NFκB translocation, the translocation of STAT1 and SMAD2/3 in response to IL-1β was also investigated. Previous studies have shown that STAT1 and SMAD2/3 do not translocate to the nucleus with either IL-1β or TNF treatment in adult human pericytes [3]. In agreement with these studies, SMAD2/3 did not exhibit nuclear translocation in HPF pericytes (Figure S5-S6). Interestingly, the iPSC-derived pericytes demonstrated STAT1 translocation at day 21 in 50.2 ± 26.3% of cells (Figure 4A-B). However, this response is absent in day 42 iPSC-derived pericytes, indicating that a STAT1 response may only be present in immature brain pericytes (Figure 4C-D). HPF pericytes also exhibited STAT1 translocation in 26.9 ± 2.59% of cells (Figure 4E-F). Our data suggests the presence of heterogeneity within the HPF pericyte population used in this study, possibly due to the presence of pericyte precursors - a consequence of the foetal origin of HPF pericytes. Additionally, these data identify important inflammatory differences between iPSC-derived brain pericytes and HPF pericytes.

**Figure 4:**
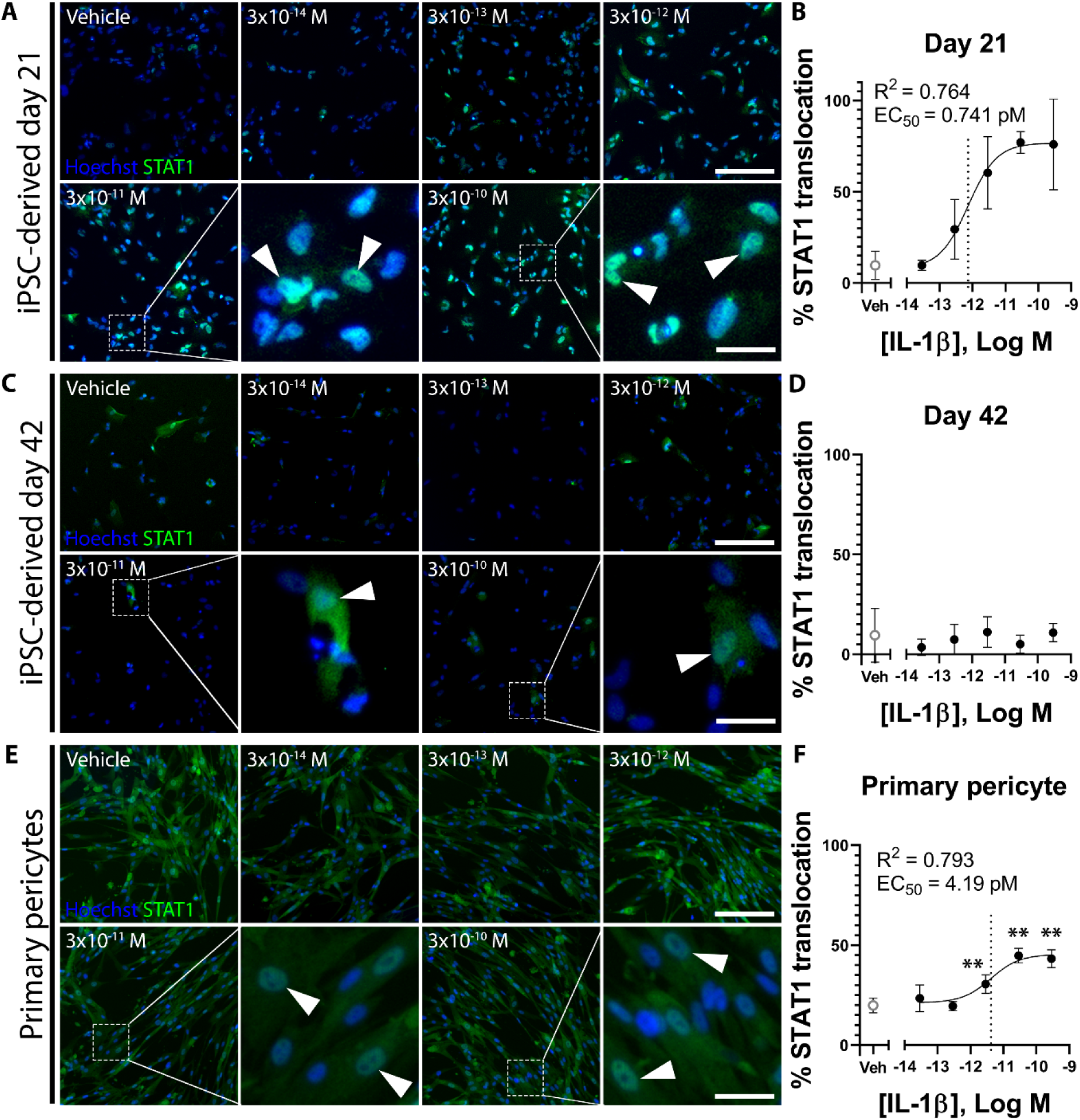
STAT1 translocates to the nucleus with IL-1β treatment in day 21 iPSC-derived pericytes and a subset of primary human pericytes, but not day 42 iPSC-derived pericytes. Immunofluorescence images demonstrating the subcellular localisation of STAT1 in response to increasing concentrations of IL-1β in day 21 iPSC-derived pericytes (A), day 42 iPSC-derived pericytes (C), and human primary foetal (HPF) brain pericytes (E). Images of iPSC-derived pericytes are quantified using an ImageJ macro to generate concentration-response curves (B, D) which demonstrate STAT1 translocation at a potent EC_50_ of 0.74pM in day 21 iPSC-derived pericytes (B, dotted line), but not in day 42 iPSC-derived pericytes. Images of HPF pericytes are quantified using MetaXpress, showing STAT1 translocation at an EC_50_ of 4.19pM (dotted line, F) in a minor subset of HPF pericytes. Data presented is one representative experiment of two experimental repeats with the iPSC-derived pericytes, and one representative experiment of three experimental repeats with the HPF pericytes (see Table S9). Scale bar = 200µm with exception of 40µm for all further magnified images. White arrows indicate nuclear STAT1. Error bars represent standard deviation. Statistical significance was determined using one-way ANOVA with Bonferroni’s multiple comparisons test. * = P<0.05, ** = P<0.01, *** = P<0.001.

The response of iPSC-derived and HPF pericytes to TNF was also investigated. It has been established that primary adult human brain pericytes exhibit nuclear translocation of NFκB in response to TNF, with the absence of STAT1 translocation [3]. However, this pathway has not been investigated in iPSC-derived pericytes. Both iPSC-derived and HPF pericytes demonstrated potent NFκB translocation in response to TNF treatment as reflected by the low EC50 values of 7.73pM in day 21 iPSC-derived pericytes, and 4.90pM in HPF pericytes (Figure 5A-F). A lack of day 42 iPSC-derived pericyte yield required the prioritisation of functional experiments, reducing the number of cells available for TNF-induced signalling characterisation. However, preliminary data for day 42 iPSC-derived pericytes showed an incomplete concentration-response curve, appearing very similar to day 21 cells between vehicle and 28.9pM, indicating that the TNF response may be conserved between the two cultures (Figure 5A-D). Of the day 21 iPSC-derived pericytes 46.1% demonstrated NFκB translocation in response to TNF, with preliminary data in day 42 iPSC-derived pericytes suggesting this increases to at least 49% (Figure 5A-D). This can be compared to the HPF pericytes where 77.2% of the cells were responsive (Figure 5E-F). While both iPSC-derived pericytes and HPF pericytes demonstrated NFκB translocation in response to TNF treatment, similar to the IL-1β response that the HPF pericytes exhibit increased responsiveness compared to iPSC-derived pericytes.

**Figure 5:**
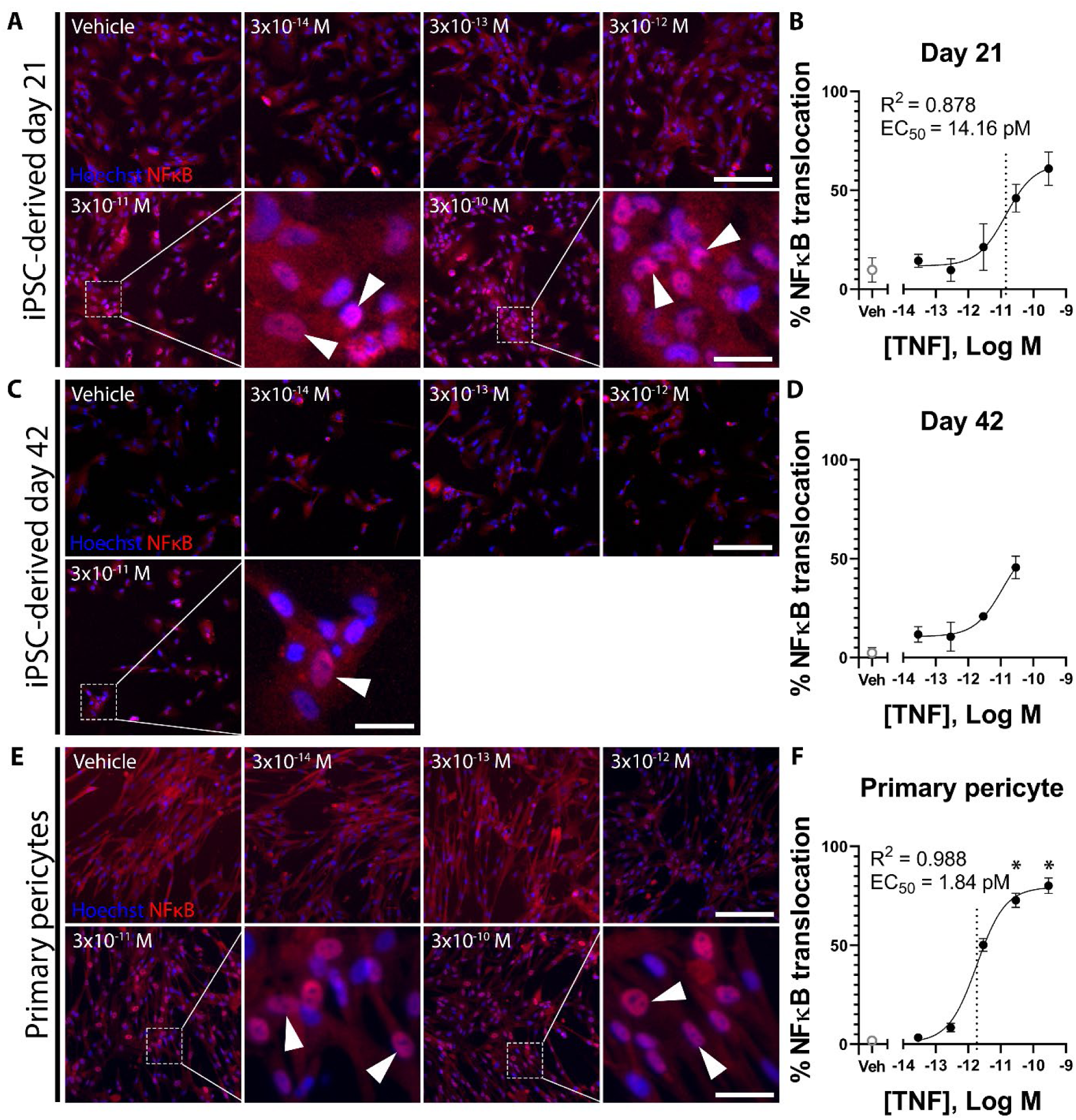
NFκB translocates to the nucleus with TNF treatment in day 21 and day 42 iPSC-derived pericytes and primary human pericytes. Immunofluorescence images demonstrating nuclear translocation of NFκB in response to increasing concentrations of TNF in day 21 iPSC-derived pericytes (A), day 42 iPSC-derived pericytes (C), and human primary foetal (HPF) brain pericytes (E). Images of iPSC-derived pericytes are quantified using an ImageJ macro to generate concentration-response curves (B, D) which demonstrates NFκB translocation at an EC_50_ of 14.2pM in day 21 iPSC-derived pericytes (B dotted line). The concentration-response curve did not plateau in day 42 iPSC-derived pericytes due to a lack of cell viability at the highest treatment concentration (D). Images of HPF pericytes are quantified using MetaXpress, showing NFκB translocation at an EC_50_ of 1.84pM (dotted line, F). Data presented is one representative experiment of two experimental repeats with the day 21 iPSC-derived pericytes, one experimental repeat with the day 42 iPSC-derived pericytes, and one representative experiment of three experimental repeats with the HPF pericytes (see Table S9). Scale bar = 200µm with exception of 40µm for all further magnified images. White arrows indicate nuclear NFκB. Error bars represent standard deviation. Statistical significance was determined using one-way ANOVA with Bonferroni’s multiple comparisons test. * = P<0.05, ** = P<0.01, *** = P<0.001.

Next, we investigated the effect of TNF on STAT1 and SMAD2/3 translocation. In line with a previously published study using primary adult human brain pericytes [3], translocation of STAT1 was not observed in either day 21 nor day 42 iPSC-derived pericytes following TNF treatment (Figure 6A-D). Interestingly, STAT1 translocated to the nucleus in 18.3% of HPF pericytes at very low concentrations, with an EC50 of 0.924pM (Figure 6E-F). This finding reveals a subset of HPF pericytes which possess a sensitive STAT1 response. Higher levels of cytoplasmic STAT1 were detected in HPF pericytes compared to iPSC-derived pericytes, indicating that HPF pericytes may express higher levels of STAT1 than iPSC-derived pericytes (Figure 6 E-F). Together, these data show differences in STAT1 activation between iPSC-derived pericytes and HPF pericytes.

**Figure 6:**
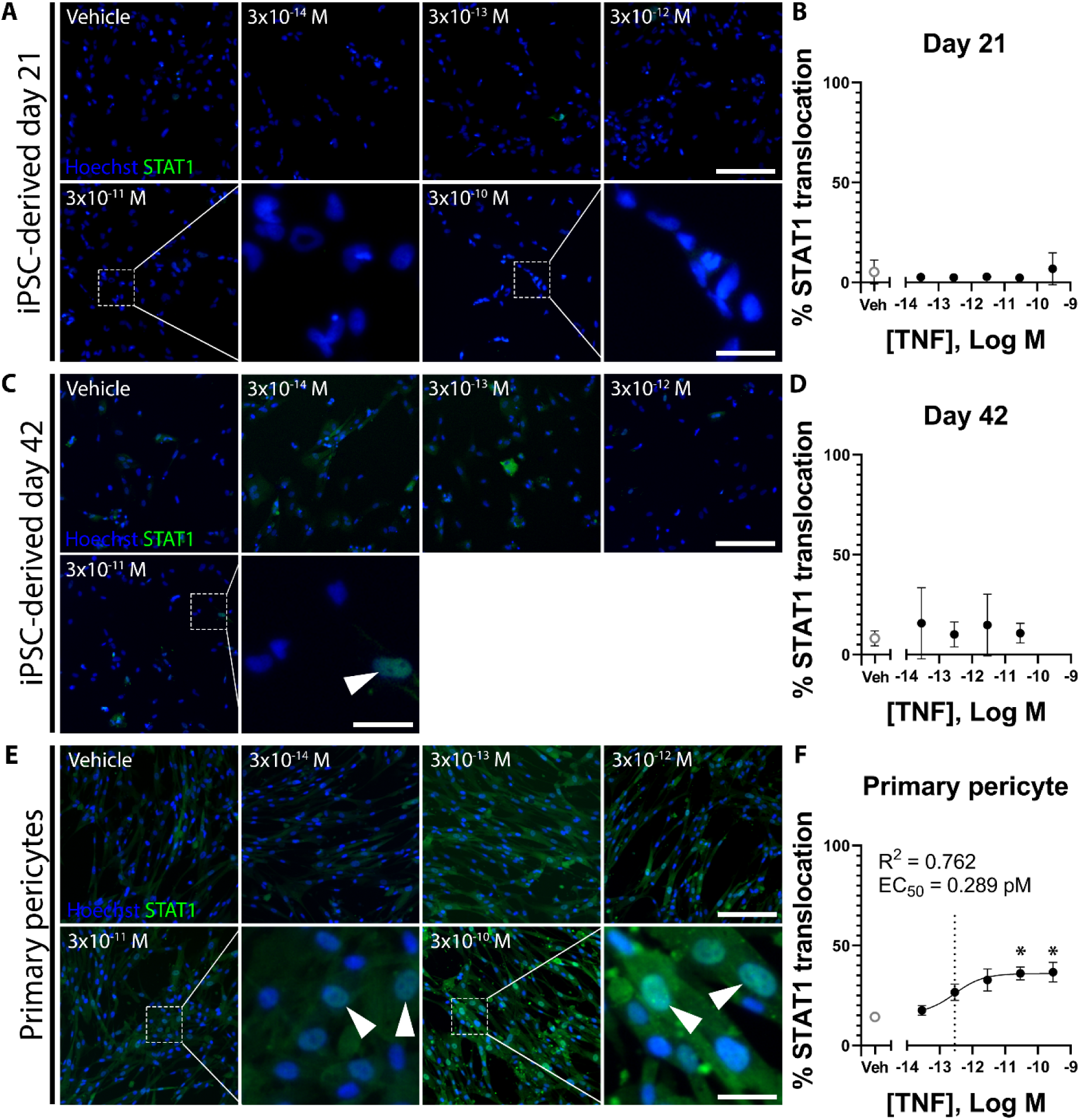
STAT1 does not translocate to the nucleus with TNF treatment in day 21 and day 42 iPSC-derived pericytes but does in a subset of primary human pericytes. Immunofluorescence images demonstrating the subcellular localisation of STAT1 in response to increasing concentrations of TNF in day 21 iPSC-derived pericytes (A), day 42 iPSC-derived pericytes (C), and human primary foetal (HPF) pericytes (E). Images of iPSC-derived pericytes are quantified using an ImageJ macro to generate concentration-response curves (B, D) which shows no STAT1 translocation in response to TNF treatment. The day 42 iPSC-derived pericytes lacked cell viability at the highest treatment concentration (C, D). Images of HPF pericytes are quantified using MetaXpress, showing very potent STAT1 translocation in a minor subset of cells at an EC_50_ of 0.289pM (dotted line, F). Data presented is one representative experiment of two experimental repeats with the day 21 iPSC-derived pericytes, one experimental repeat with the day 42 iPSC-derived pericytes, and one representative experiment of three experimental repeats with the HPF pericytes (see Table S9). Scale bar = 200µm with exception of 40µm for all further magnified images. White arrows indicate nuclear STAT1. Error bars represent standard deviation. Statistical significance was determined using one-way ANOVA with Bonferroni’s multiple comparisons test. * = P<0.05, ** = P<0.01, *** = P<0.001.

### iPSC-derived brain pericytes exhibit different secretory and phagocytic responses to inflammatory cytokine treatment

The IL-1β- and TNF-induced secretory responses in primary adult human brain pericytes have been well studied. Brain pericytes have been shown to secrete a plethora of inflammatory cytokines and chemokines when stimulated with either IL-1β or TNF, further propagating the inflammatory environment. After demonstrating that iPSC-derived pericytes exhibit NFκB translocation in response to IL-1β and TNF, the secretion of chemokines, cytokines, adhesion molecules, and growth factors in response to IL-1β and TNF was investigated to see whether iPSC-derived pericytes exhibit the same secretory profile as that of HPF pericytes (Figure 7, Figure 8).

**Figure 7:**
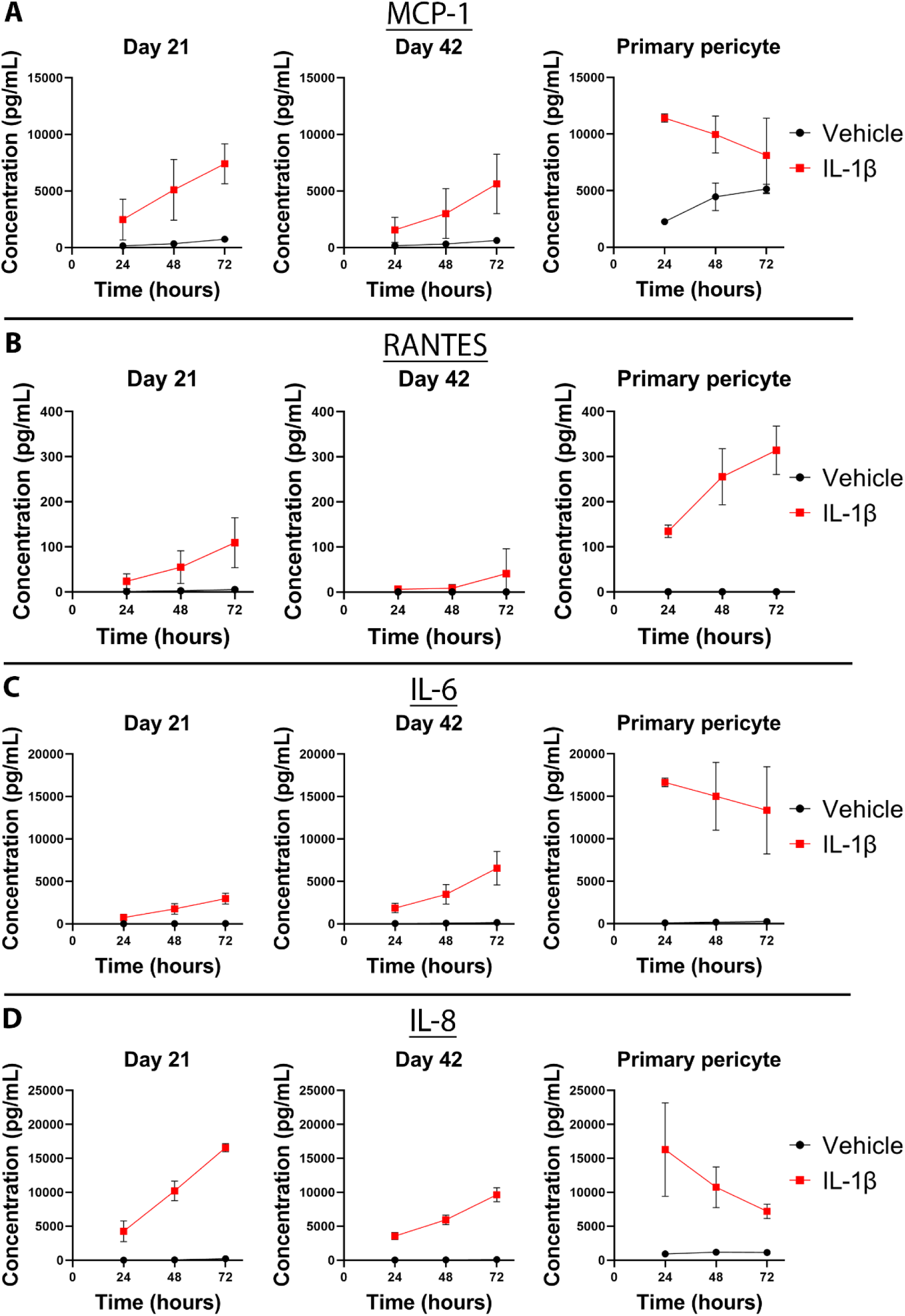
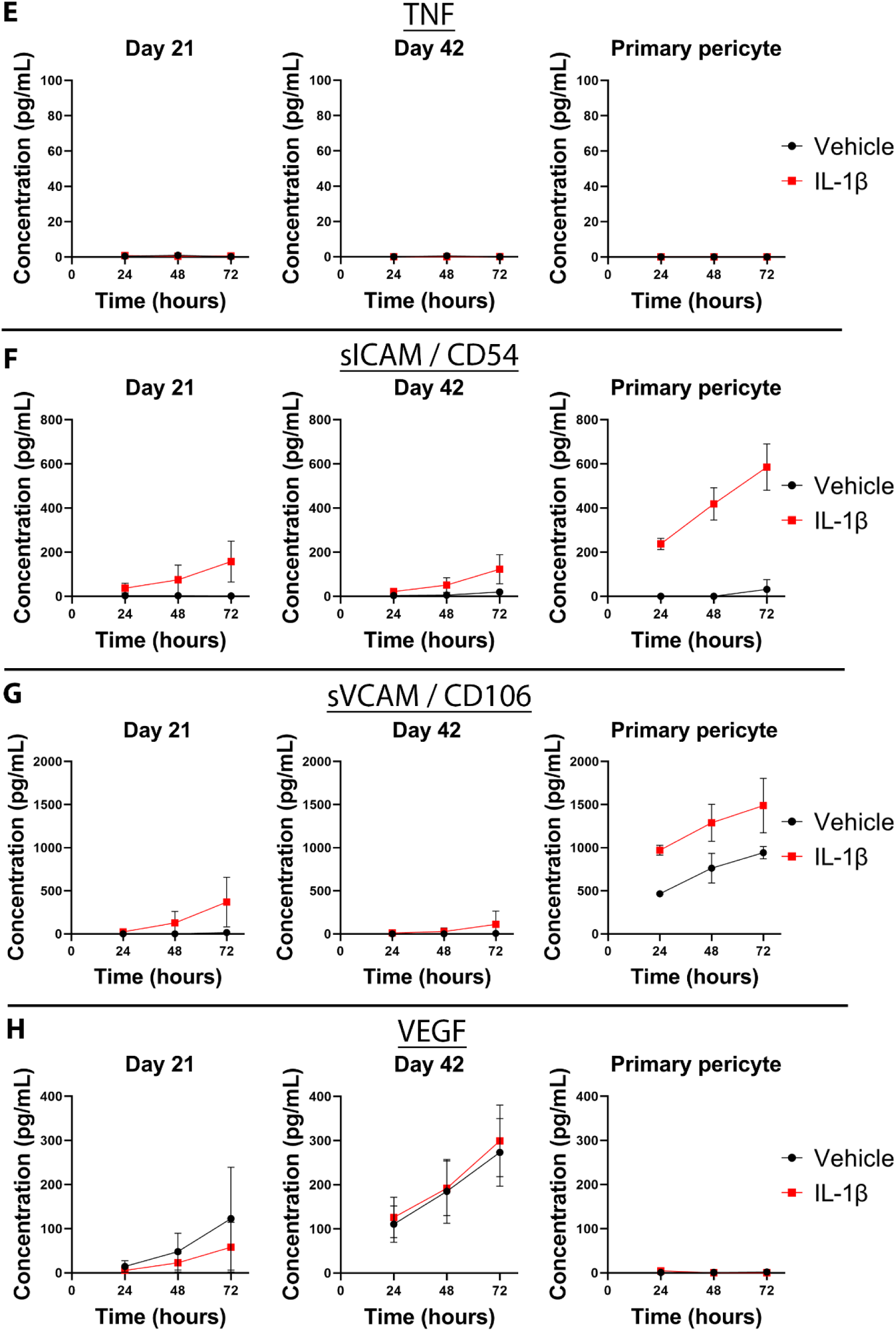
IL-1β induces distinct inflammatory cytokine and chemokine secretion in day 21 iPSC-derived pericytes, day 42 iPSC-derived pericytes, and primary human pericytes. Graphs demonstrating secretion of (A) MCP-1, (B) RANTES, (C) IL-6, (D) IL-8, (E) TNF, (F) sICAM, (G) sVCAM, and (H) VEGF in response to IL-1β treatment (28.9pM or 500pg/mL) or vehicle treatment at 24-, 48-, and 72-hours following treatment. Data presented is averaged from two experimental repeats. Error bars represent standard deviation.

**Figure 8:**
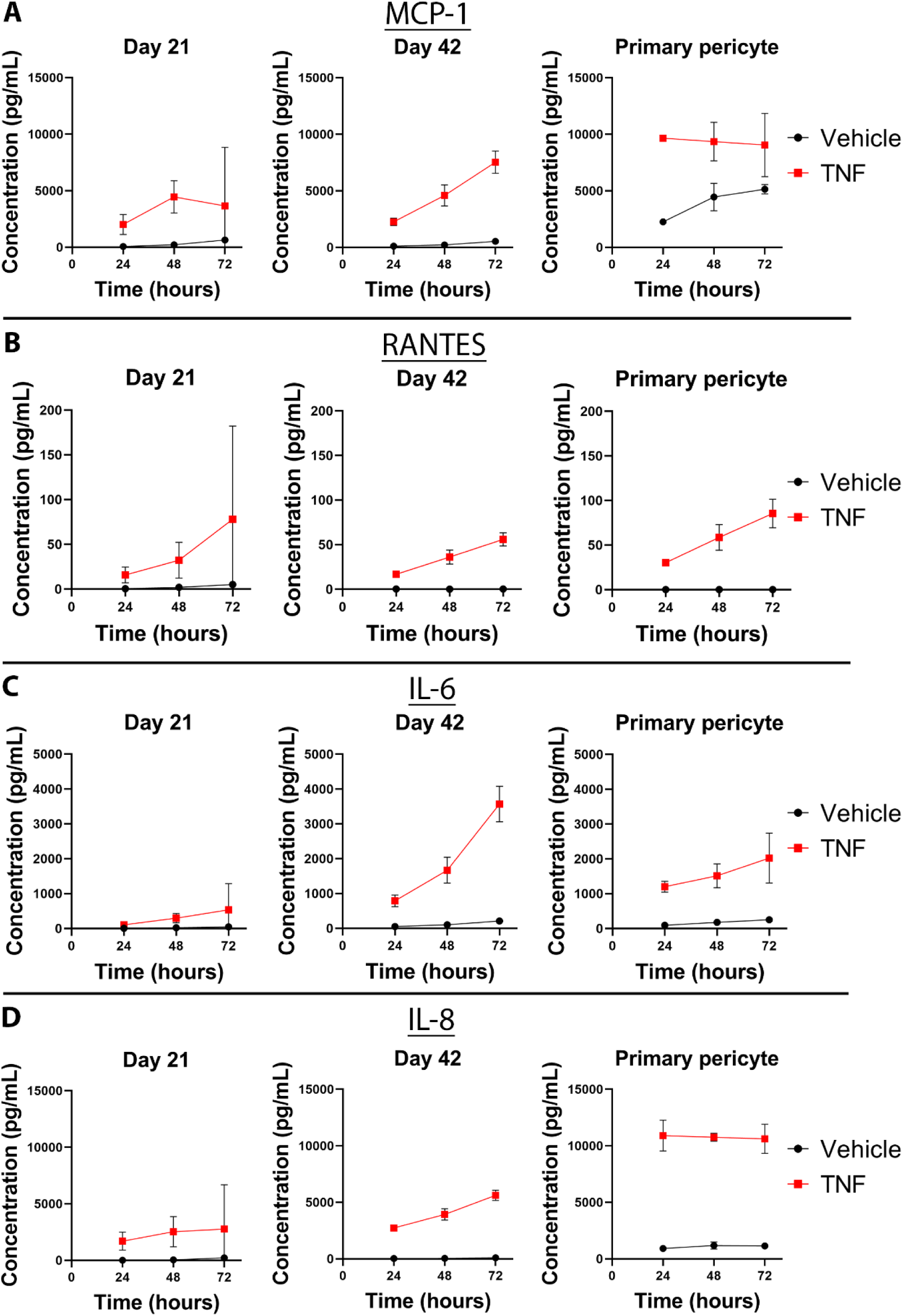
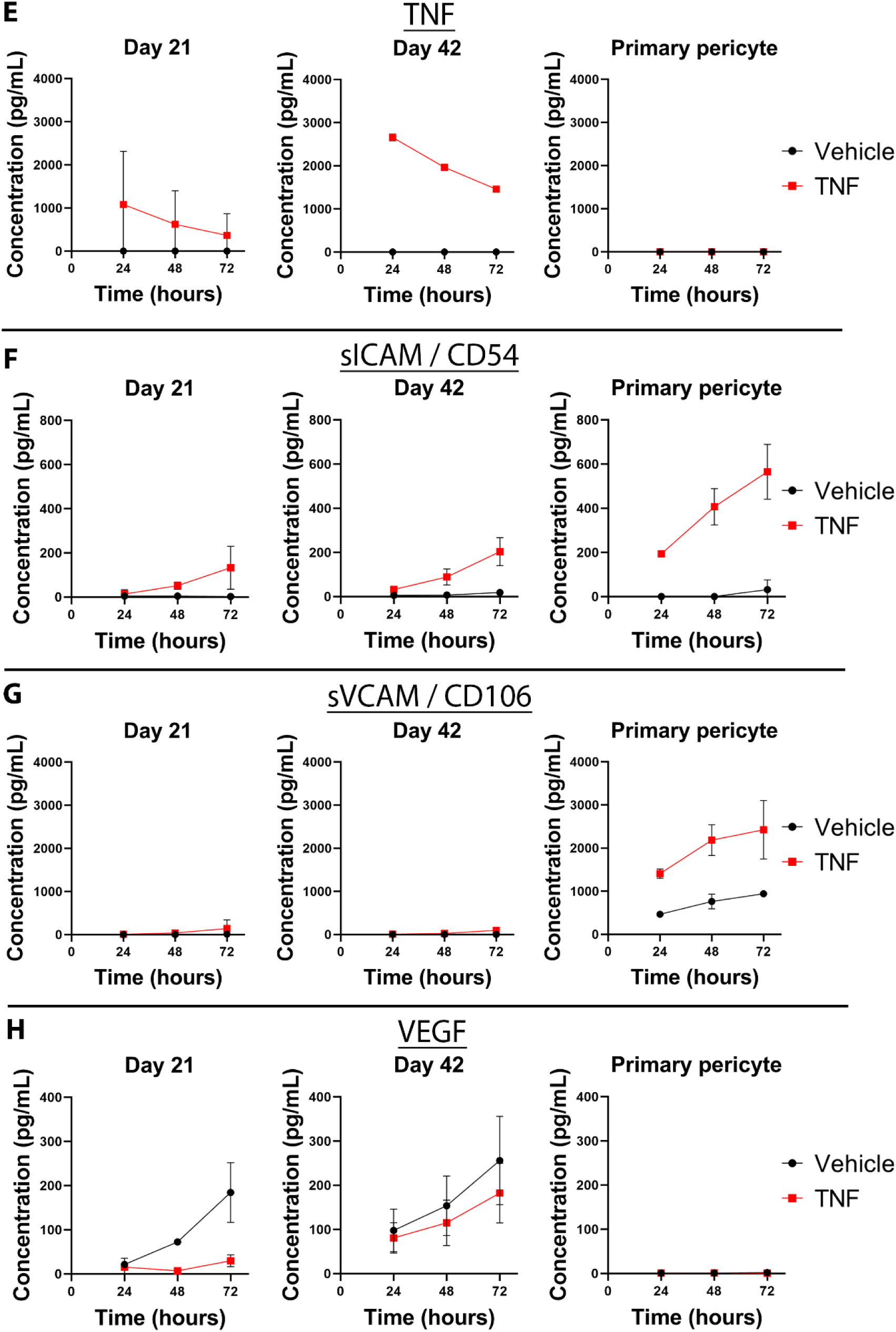
TNF induces distinct inflammatory cytokine and chemokine secretion in day 21 iPSC-derived pericytes, day 42 iPSC-derived pericytes, and primary human pericytes. Graphs demonstrating secretion of (A) MCP-1, (B) RANTES, (C) IL-6, (D) IL-8, (E) TNF, (F) sICAM, (G) sVCAM, and (H) VEGF in response to TNF treatment (28.9pM or 500pg/mL) or vehicle treatment at 24-, 48-, and 72-hours following treatment. Data presented is averaged from two experimental repeats.

The concentration of secreted chemokines MCP-1, fractalkine, and RANTES was investigated because adult human pericytes have previously been found to secrete all these factors in response to IL-1β and TNF treatment [3]. In both day 21 and day 42 iPSC-derived pericytes, MCP-1 was detected at increasing concentrations over time, suggesting the consistent secretion of MCP-1 across the 72 hours (Figure 7A). In contrast, the concentration of MCP-1 in HPF pericyte cultures was highest after 24 hours and decreased after this timepoint, indicative of regulatory mechanisms to control local MCP-1 activity following IL-1β or TNF treatment (Figure 7A, Figure 8A). Fractalkine was detected in cultures of day 21 iPSC-derived pericytes after 48 hours with IL-1β treatment, with a higher concentration observed after 72 hours (Figure 7B). Fractalkine was only detected after 72 hours following TNF treatment (Figure 8B). This suggests that after a delay, fractalkine is consistently secreted in these cultures, with a greater delay in expression following TNF treatment. Curiously, this response was not present in day 42 iPSC-derived pericytes, which lacked any fractalkine secretion with either IL-1β or TNF treatment across the investigated 72 hours (Figure 7B, Figure 8B). The concentration of fractalkine was highest after 24 hours but was maintained across the following 72 hours in IL-1β- and TNF-treated HPF pericyte cultures (Figure 7B, Figure 8B). This could be due to downregulation of fractalkine secretion after the initial 24 hours. Induction of RANTES secretion was observed with both IL-1β and TNF treatment in day 21 iPSC-derived pericytes and HPF pericytes, with the observed concentration of RANTES increasing over time, indicative of consistent secretion across this period (Figure 7C, Figure 8C). RANTES was not observed in the medium of IL-1β-treated day 42 iPSC-derived pericytes across the first 48 hours, though a small amount was detected after 72 hours of IL-1β treatment, suggesting that the culture may contain a subset of cells which secrete RANTES (Figure 7C). RANTES was observed with TNF treatment at comparable concentrations to HPF pericytes, with higher concentrations observed over time (Figure 8C).

After recruiting immune cells through chemokine secretion, adult brain pericytes can propagate the initial inflammatory signal through the production and secretion of inflammatory cytokines such as IL-6, IL-8, or TNF [3,18,19]. Both day 21 and day 42 iPSC-derived pericytes demonstrated increasing concentrations of IL-6 and IL-8 over 72 hours with both IL-1β and TNF treatment, again likely due to consistent secretion in response to treatment (Figure 7D-E, Figure 8D-E).

The highest concentration of IL-6 and IL-8 in IL-1β-treated HPF pericytes was observed after 24 hours, decreasing with time, demonstrating mechanisms to regulate IL-6 and IL-8 availability after 24 hours (Figure 7D-E). Interestingly, this response was not apparent with TNF treatment, where the concentration of IL-6 increased with time and IL-8 was maintained across the 72 hours (Figure 8D-E). This indicates the presence of different regulatory responses to IL-1β and TNF treatment. Secreted TNF was not detected with IL-1β treatment, suggesting its production is not induced with IL-1β treatment in either iPSC-derived pericytes or HPF pericytes (Figure 7F). TNF was observed in the media of TNF-treated day 21 and day 42 iPSC-derived pericytes (Figure 8F). The TNF concentration decreased over time, indicating that these cells possess mechanisms to regulate TNF availability. Curiously, TNF was not observed in the medium of TNF-treated HPF pericytes (Figure 8F). This could be due to the expression of proteins which degrade or sequester available TNF within the first 24 hours, a novel function of HPF pericytes. This degradation of TNF was dependent on the presence of pericytes as TNF in medium without cells was observed after 72 hours (Figure S7).

Previous studies have found that adult brain pericytes upregulate release of soluble ICAM and sVCAM (sICAM/sVCAM)in response to IL-1β or TNF treatment, which is evidence of cleavage of membranous ICAM and VCAM [3]. Here, secretion of sICAM was induced with both IL-1β and TNF treatment in day 21 iPSC-derived pericytes, day 42 iPSC-derived pericytes, and HPF pericytes, with increasing concentrations of sICAM detected over time (Figure 7G, Figure 8G). This response in iPSC-derived pericytes was negligible after 24 hours, requiring 48 hours for sICAM to be observed in the medium (Figure 7G, Figure 8G). HPF pericytes demonstrated robust secretion of sICAM, with higher concentrations of sICAM observed across all time points when compared with the iPSC-derived pericytes (Figure 7G, Figure 8G). The day 21 and day 42 iPSC-derived pericytes demonstrated a similar sVCAM secretory responses to IL-1β and TNF treatment, with a modest increase in sVCAM concentration observed in the medium of both cultures over time (Figure 7H, Figure 7H). Again, the day 21 iPSC-derived pericytes did not show a difference to vehicle in the first 24 hours, requiring at least 48 hours before sVCAM was observed in the medium (Figure 7G). The day 42 iPSC-derived pericyte cultures required more time for sVCAM to be observed in the medium, with differences between IL-1β-, and vehicle-treated samples only seen after 72 hours (Figure 7H, Figure 8H). Higher concentrations of sVCAM were observed in the HPF pericyte cell medium (Figure 7H). Interestingly, higher concentrations of sVCAM were observed in vehicle-treated HPF pericyte medium, suggesting that sVCAM is released under basal conditions (Figure 7H). This is corroborated by the increasing concentrations of sVCAM observed over time in these cultures. Treatment with IL-1β or TNF increases the concentration of sVCAM observed across all timepoints but does not change the rate of sVCAM secretion after 24 hours (Figure 7G). It is likely that the effect of IL-1β and TNF treatment on sVCAM secretion occurs over the initial 24 hours but does not change sVCAM secretion after this timepoint.

VEGF was not observed in treated HPF pericyte medium across all time points (Figure 7I, Figure 8I). However, VEGF was observed in day 21 and day 42 iPSC-derived pericyte cultures (Figure 7I, Figure 8I). The concentration of VEGF increased with time in vehicle-treated samples, suggesting that VEGF is secreted at rest by both day 21 and day 42 iPSC-derived pericytes (Figure 7I, Figure 8I). This secretion was not changed by IL-1β treatment but was reduced with TNF treatment in day 21 iPSC-derived pericytes. Collectively, these data demonstrate many differences between HPF and iPSC-derived pericyte secretory responses to IL-1β.

Another important function of adult brain pericytes is their ability to phagocytose material that passes through the blood-brain barrier [4,5,20]. To compare the phagocytic activity of our iPSC-derived pericytes with HPF pericytes, we used two different phagocytosis assays utilizing fluorescent beads combined with either fluorescent microscopy or flow cytometry. Phagocytic cells could be identified using fluorescent microscopy by clumps of fluorescent bead associating with single nuclei (Figure 9A, white arrows). Fewer beads associated with HPF pericytes compared to iPSC-derived pericytes (Figure 9A). Fluorescent microscopy did not reveal any changes in the number of beads per cell when either HPF pericytes or iPSC-derived pericytes were treated with IL-1β or TNF (Figure 9A). Some cells appeared to lack phagocytic ability in both the HPF pericyte and iPSC-derived pericyte cell cultures (Figure 9A).

**Figure 9:**
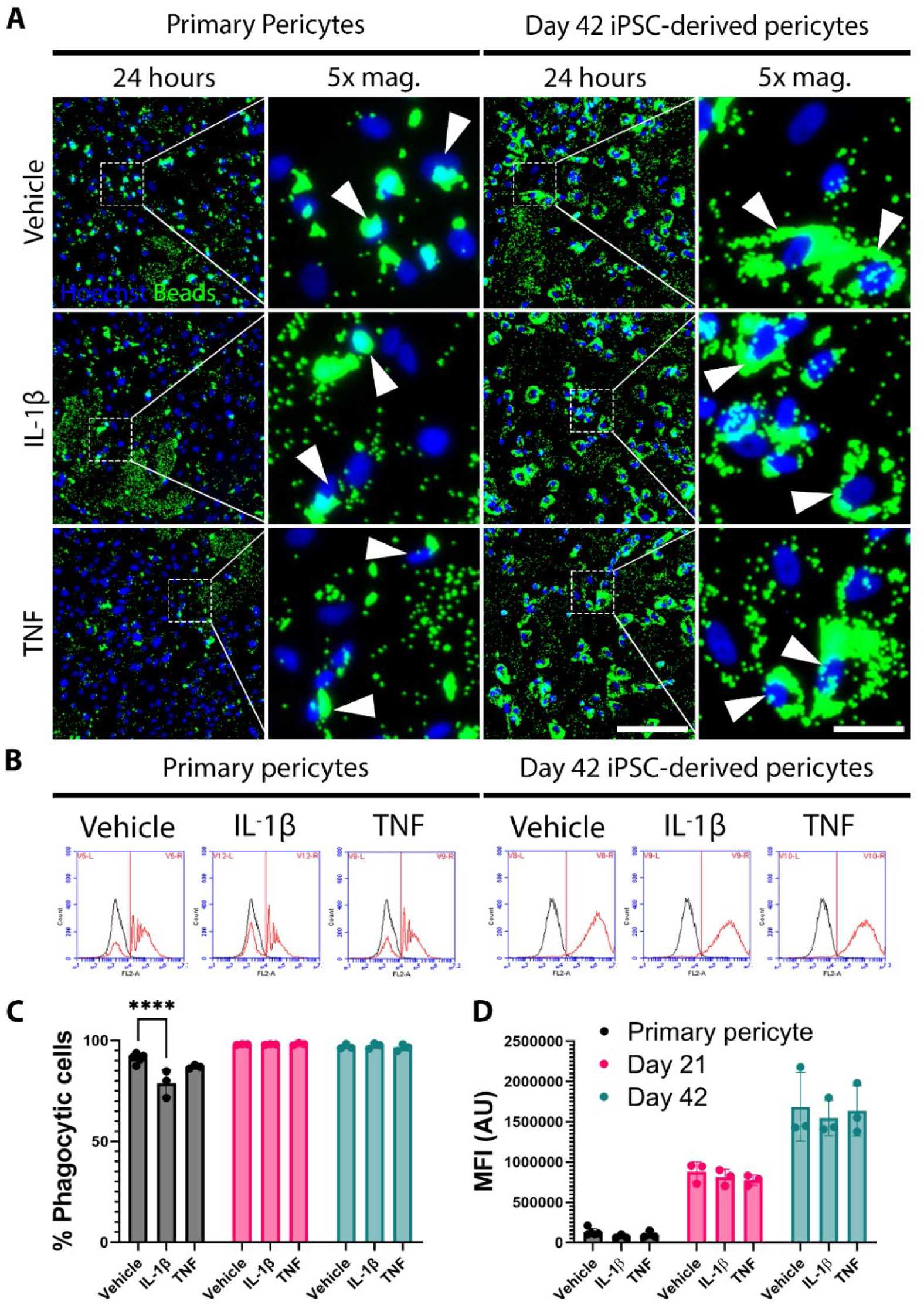
iPSC-derived pericytes exhibit increased rates of phagocytosis compared to primary human pericytes. (A) Immunofluorescence images comparing the abundance of phagocytosed fluorescent beads in human primary foetal (HPF) pericytes (left) and iPSC-derived pericytes (right). (B) Flow cytometry histo-plots show cultured primary cells to contain phagocytic (red) and non-phagocytic (black) cells. The auto-fluorescent threshold is denoted by the vertical red line. The gating strategy for flow cytometric analysis can be found in Figure S1. (C,D) Quantification of histo-plots shows a significant reduction in percentage of phagocytic HPF pericytes with IL-1β treatment, but no change in either day 21 or 42 iPSC-derived pericytes. No change in mean fluorescent intensity (MFI) was observed with either IL-1β or TNF treatment, though day 21 and day 42 iPSC-derived pericytes exhibited more phagocytic activity than HPF pericytes (D). Quantitative data presented is averaged from three to five experimental repeats. Significance is determined using a 2-way ANOVA with Tukey’s multiple comparisons test.

Assessment of phagocytosis by flow cytometry corroborated these findings, demonstrating the iPSC-derived pericytes phagocytosed more fluorescent beads (Figure 9B-D). Additionally, as flow cytometry allows for interrogation of individual cell fluorescence, the percentage of phagocytic cells in each cell population could be assessed. After vehicle treatment, 91.1% of HPF pericytes exhibited phagocytic activity, compared to 98.0% of day 21 iPSC-derived pericytes and 96.9% of day 42 iPSC-derived pericytes (Figure 9C-D). Importantly, HPF pericytes display lower rates of phagocytosis to day 21 iPSC-derived pericytes, which show lower phagocytosis than day 42 iPSC-derived pericytes as measured by mean fluorescent intensity (MFI) (Figure 9D). When treated with IL-1β, HPF pericytes exhibited a significant (p<0.0001) decrease in the percentage of phagocytic cells, but not the MFI of phagocytic cells (Figure 9B-D). The iPSC-derived pericytes did not exhibit any changes in phagocytosis when treated with either IL-1β or TNF. These data show large differences in phagocytosis between HPF pericytes and iPSC-derived pericytes.

## Discussion

The use of iPSC-derived cell models is a fast-growing field that is producing promising results. Surprisingly, our study shows that the iPSC-derived pericytes we generated demonstrated a different inflammatory response to HPF pericytes and exhibited critical differences in expression and maturation compared to other studies using the same protocol [14]. Additionally, significant variation was observed between cells generated by separate differentiations. This work highlights important challenges currently faced by the field of iPSC-derived research, raising concern for researchers who may be tempted to use this technology when it is not always fit for purpose.

We generated brain pericyte-like cells in 42 days according to the published protocol by Stebbins et al. and investigated these cells at day 21 and day 42 of differentiation. Differences in several signalling pathways were observed between the response of iPSC-derived pericytes and HPF pericytes to inflammatory stimuli (Table 1). IL-1β and TNF treatment resulted in STAT1 translocation in a minor subset of HPF pericytes, a response that was present in a large proportion of day 21 iPSC-derived pericytes but absent by day 42 of differentiation. This suggests that immature pericytes exhibit a STAT1 response to IL-1β treatment. The STAT1 translocation observed in our HPF pericytes is likely the result of the foetal origin of these cells, given that this response is absent in primary adult human pericytes [3]. This immature pericyte subpopulation may also contribute to the high concentration of OCT3/4 mRNA detected in HPF pericytes. These results reveal important differences between the actions and expression profiles of adult and foetal human brain pericytes. This is a considerable limitation to the use of HPF pericytes, which must be further understood as HPF pericytes are the only human brain pericytes that are commercially available.

**Table 1:** Summary of functional differences between iPSC-derived pericytes, HPF pericytes and human primary adult pericytes.

| IL-1β Treatment | | | | | | | |
| --- | --- | --- | --- | --- | --- | --- | --- |
|  | Day 21 |  | Day 42 |  | HPF Pericytes |  | HPA Pericytes |
| NFκB translocation | +++ | | +++ | | +++ | | +++ |
| STAT1 translocation | ++ |  | 0 |  | + |  | 0 |
|  | 24h | 72h | 24h | 72h | 24h | 72h | 24h |
| MCP-1 | + | +++ | + | +++ | +++ | + | +++ |
| RANTES | 0 | ++ | 0 | 0 | ++ | +++ | + |
| IL-6 | 0 | + | 0 | ++ | +++ | +++ | +++ |
| IL-8 | + | +++ | + | ++ | +++ | ++ | +++ |
| TNF | 0 | 0 | 0 | 0 | 0 | 0 | n/a |
| sICAM | 0 | + | 0 | + | ++ | +++ | ++ |
| sVCAM | 0 | + | 0 | 0 | ++ | ++ | +++ |
| VEGF | 0 | 0 | 0 | 0 | 0 | 0 | n/a |
| TNF Treatment |  |  |  |  |  |  |  |
|  | Day 21 |  | Day 42 |  | HPF Pericytes |  | HPA Pericytes |
| NFκB translocation | ++ | | ++ | | +++ | | +++ |
| STAT1 translocation | 0 |  | 0 |  | + |  | 0 |
|  | 24h | 72h | 24h | 72h | 24h | 72h | 24h |
| MCP-1 | + | + | + | ++ | +++ | + | +++ |
| RANTES | + | ++ | + | ++ | + | ++ | + |
| IL-6 | 0 | 0 | + | +++ | + | ++ | +++ |
| IL-8 | + | + | + | ++ | +++ | +++ | +++ |
| TNF | + | 0 | +++ | ++ | 0 | 0 | n/a |
| sICAM | 0 | + | 0 | + | ++ | +++ | ++ |
| sVCAM | 0 | 0 | 0 | 0 | + | ++ | +++ |
| VEGF | 0 | -- | 0 | - | 0 | 0 | n/a |
| Basal conditions |  |  |  |  |  |  |  |
|  | Day 21 |  | Day 42 |  | HPF Pericytes |  | HPA Pericytes |
| Phagocytosis | ++ |  | +++ |  | + |  | + |

0 indicates no response. +/++/+++ indicates the intensity of an induced response. -/-- indicates the intensity of an inhibitory response. Human primary adult (HPA) pericyte data taken from: <sup>[3,4]</sup>. n/a indicates that no relevant experimental data exists.

Changes were also observed in treatment sensitivity, with IL-1β and TNF treatment causing a potent and robust NFκB response in HPF pericytes and a weaker and less consistent response in iPSC-derived pericytes. It should be noted that the HPF pericytes responded to IL-1β and TNF at concentrations 100-fold lower than comparable published treatment concentrations [3,16,20]. The HPF pericytes used here may be more sensitive than the primary adult human pericytes used in previous studies, or the generally accepted concentration of IL-1β and TNF treatment in brain pericytes is too high. Care should be taken to ensure that the responses seen with more concentrated treatments of IL-1β and TNF are due to canonical responses, and not non-specific signalling.

Substantial differences were also observed between the inflammatory secretome of iPSC-derived pericytes and HPF pericytes (Table 1). IL-1β treatment resulted in acute secretion of MCP-1, fractalkine, IL-6, and IL-8 in primary cells, with the concentrations of these secreted factors decreasing over time, indicating the presence of regulatory mechanisms to control the levels of these factors over time. These regulatory mechanisms are likely to be absent in iPSC-derived pericytes, where the concentrations of MCP-1, fractalkine, IL-6, and IL-8 increase over time. Higher concentrations of MCP-1, RANTES, IL-6, and ICAM were also observed in IL-1β treated HPF pericytes, perhaps due to the increased sensitivity of these cells to inflammatory stimuli. Finally, iPSC-derived pericytes exhibited much higher rates of phagocytosis than HPF pericytes, which wasn’t changed when treated with common inflammatory factors IL-1β or TNF. Conversely, HPF pericytes exhibited a decrease in observed phagocytosis when treated with IL-1β or TNF. Collectively, these results demonstrate many differences between the signalling and secretory responses of iPSC-derived pericytes and HPF pericytes, providing uncertainty as to which cell type best models the inflammatory actions of adult brain pericytes *in vivo*.

As well as differences in inflammatory response, the iPSC-derived pericyte protocol suffered from poor reproducibility, generating cells with variable responses and inconsistent cell yield. A 42-day differentiation protocol to produce mature cells is an enormous investment of time and money and given the lack of consistency demonstrated by these cells, significant financial and experimental redundancy is necessary to generate meaningful results. This makes iPSC-derived models difficult to set-up in lab groups that have limited resources. The cells generated by this protocol exhibited differences to the reported characterisation of the original protocol by Stebbins and colleagues [14]. The original protocol generated pericyte-like cells by day 21, while in our hands pericyte-like cells were only observed after 42 days. Notably, αSMA was absent in the original protocol and abundant in our iPSC-derived pericytes. This lab-specific variation in the differentiated cells could stem from the use of different iPSC sources. Differences in cell passage number, growth factor suppliers, and specific media formulation could be responsible for the significant differences between the generated iPSC-derived pericytes, necessitating protocol optimisation before use.

Despite the plethora of markers that are associated with brain pericytes, a brain pericyte-specific marker has yet to be found. The lack of a defining brain pericyte marker may be hindering their use as an iPSC-derived model due to the presence of currently undefined brain pericyte subtypes. Similar limitations are found with mesenchymal stem cells, which also lack a cell-specific marker. Interestingly, differences have also recently been reported between iPSC-derived mesenchymal stem cells and primary mesenchymal stem cells [21,22]. Perhaps the incomplete characterisation of brain pericytes and mesenchymal stem cells contributes to the variable results demonstrated by different lab groups. Further characterisation of brain pericytes is required such that we can be more certain that these iPSC-derived cells exhibit a true brain pericyte phenotype.

Many protocols have claimed to differentiate pericytes from iPSCs, showing cells which express several pericyte markers such as PDGFRβ, NG2, or αSMA (see Table S1). However, this is not sufficient to distinguish pericytes from either fibroblasts or smooth muscle cells. The potassium channel subunit *ABCC9* is expressed in pericytes, but is absent in smooth muscle cells, and may help distinguish these cell types [23,24]. However, a marker is yet to be found which comprehensively distinguishes fibroblasts from pericytes. We saw an increase in expression of *ABCC9* across NCSC priming (days 1-15), but a reduction across pericyte differentiation (days 15-42). This may be due to the cells acquiring a more smooth muscle-like phenotype over time *in vitro*, a phenomenon that has been reported previously in primary adult human brain pericytes [4]. Of the previous pericyte differentiation protocols, the majority use a mesodermal differentiation protocol likely to produce pericytes of other organs of the body (Table S1) [1]. We used a neural crest differentiation protocol to generate brain pericytes and assessed this through the expression of brain pericyte markers *FOXF2*, *FOXC1*, and *VTN*. Expression of these markers is found in other tissues of the body but not other pericytes, with recent work showing that the expression of these markers is critical in brain pericytes to contribute to BBB function [8,11,25]. The presence of *FOXF2* and *FOXC1* in the iPSC-derived pericytes generated here provides evidence of a brain pericyte phenotype. Expression of *VTN* as a brain pericyte marker came from an RNA-Seq study from whole mouse brain, followed by studies demonstrating that pericyte-derived vitronectin reduces transcytosis in brain endothelial cells [8,26]. However, a more recent RNA-Seq study comparing human and mouse microvascular cell expression has shown that *VTN* may be specifically expressed in mouse brain pericytes and not human brain pericytes [7]. The work presented here shows *VTN* to be expressed in both iPSC-derived pericytes and HPF pericytes, suggesting that there are states in which human brain pericytes do express *VTN*. While additional research is required to confirm the presence and function of *VTN* in human brain pericytes, investigation of FOXF2 and FOXC1 should be standard for studies claiming to work with iPSC-derived brain pericytes.

Interestingly, TNF-treated HPF pericytes degraded or sequestered TNF such that none was observed in the culture medium after 24 hours. This may have implications on the neuroinflammatory response of adult human brain pericytes and introduces a novel role of brain pericytes as a neuroinflammatory quencher. The iPSC-derived pericytes also exhibited the ability to degrade or sequester TNF but was less efficient than the HPF pericytes. HPF pericytes respond to TNF, indicating the presence of TNF receptors (TNFRI/TNFRII) on their cell surface [3]. Perhaps the internalisation and degradation of these receptors acts as a mechanism to remove TNF from the environment. Alternatively, mouse brain pericytes have been shown to secrete matrixmetalloprotease-2 (MMP-2) and MMP-9, offering potential routes of TNF degradation [27,28]. It is likely that this is a function of brain pericytes to soak up inflammatory cytokines, thereby protecting the integrity of the BBB. It is important that future research elucidates the mechanisms of TNF degradation as the ability to quench chronic neuroinflammation may provide therapeutic benefit for many neurodegenerative diseases.

## Conclusions

With time, cell differentiation protocols will be optimised to reduce the variability of iPSC-derived cells, allowing for comparisons to be drawn between labs using similar or identical protocols. In the meantime, it is important that the limitations of iPSC-derived cell models are noted. Most current iPSC-derived cell models do not discuss all the challenges involved with this work, enticing other lab groups to investigate the use of iPSC-derived models in their labs. If the limitations of iPSC-derived cell models are more openly shared, then researchers will be able to make better informed decisions regarding whether this system is right for them. For now, labs adopting iPSC-derived cell culture methods should be prepared to perform their own characterisation of the produced cells and expect to see differences with other protocols in the field. We demonstrated numerous functional differences between iPSC-derived pericytes and HPF pericytes. Future work is required to increase the consistency of the iPSC protocol and to uncover which human brain pericyte phenotype best represents that of adult brain pericytes *in vivo*.

## Declarations

### Author contributions

**Samuel McCullough**: Conceptulization, Data curation, Formal analysis, Investigation, Methodology, Writing – original draft, Writing – review and editing. **Eliene Albers**: Conceptualization, Formal analysis, Supervision, Writing – review & editing. **Akshata Anchan**: Investigation, Methodology, Supervision, Writing – review & editing. **Jane Yu**: Methodology, Supervision. **Simon O’Carroll**: Resources, Supervision, Writing – review & editing. **Bronwen Connor**: Conceptualization, Formal analysis, Methodology, Supervision, Writing – review & editing. **E. Scott Graham**: Conceptualization, Formal analysis, Funding acquisition, Methodology, Project administration, Resources, Software, Supervision, Writing – review & editing

## Acknowledgements

This work was supported by the Ministry of Business, Innovation and Employment, New Zealand and the Department of Anatomy and Medical Imaging, University of Auckland. None of these funding bodies had any role in the research or interpretation of the data. Jacqueline Ross developed the macro used to quantify the concentration-response images from iPSC-derived pericytes. Diagrams were generated using BioRender.

## Conflict of interest statement

The authors declare that they have no conflicts of interest.

## Ethics statement

Ethics approval was not needed in this study.

## Data availability statement

All data used to support the findings of this study is included within the article. Raw data used to generate the figures are available from the corresponding author upon request.

## Notes

### Competing Interest Statement

The authors have declared no competing interest.

