## Supplementary material for "Human iPSC-derived brain pericytes exhibit differences in inflammatory activation compared to primary human brain pericytes"

### Supplemental material

**Table S1: Summary of protocols for iPSC-derived pericyte differentiation.**

| Authors | Year first published | Number of Citations | Germ line of derived cells | Notes |
|---|---|---|---|---|
| Taura et al. | 2009 | 160 | Mesoderm | Uses feeder cells. Analyses cells through FACS. |
| Dar et al. | 2012 | 175 | Mesoderm | Generates pericytes and endothelial cells. Analyses cells through FACS. |
| Kusuma et al. | 2013 | 172 | Mesoderm | Isolates pericyte-like cells through growth advantages. |
| Masumoto et al. | 2014 | 201 | Mesoderm | Generates a cardiac tissue sheet. |
| Orlova et al. | 2014 | 226 | Mesoderm | Generates pericytes and endothelial cells. Isolates pericyte-like cells through Dynabead cell sorting. |
| Park et al. | 2014 | 66 | Mesoderm | Isolates pericyte-like cells through FACS. |
| Ren et al. | 2015 | 166 | Mesoderm | Isolates pericyte-like cells through FACS. |
| Kumar et al. | 2017 | 117 | Mesoderm | Isolates pericyte-like cells through growth advantages. |
| Maffioletti et al. | 2018 | 166 | Unknown | Generate a muscle organoid. |
| Faal et al. | 2019 | 41 | Mesoderm | Isolates pericyte-like cells through growth advantages. |
| Wimmer et al. | 2019 | 71 | Mesoderm | Generates a vascular organoid. |
| **Faal et al.** | **2019** | **41** | **Neural Crest** | **Isolates pericyte-like cells through growth advantages.** |
| **Kelleher et al.** | **2019** | **21** | **Neural Crest** | **Isolates pericyte-like cells through growth advantages.** |
| **Stebbins et al.** | **2019** | **95** | **Neural Crest** | **Isolates pericyte-like cells through MACS.** |
| Szepes et al. | 2020 | 8 | Mesoderm | A modified version of the Orlova et al., 2014 protocol. Isolates pericyte-like cells through FACS. |

**Table S2: E6-CSFD components.**

| E6-CSFD | Concentration | Company | Catalogue# |
|---|---|---|---|
| Essential 6 | | Gibco | A1516401 |
| CHIR99021 | 1μM | Tocris | RDS442350 |
| SB431542 | 10μM | Merck | S4317 |
| FGF2 | 10ng/mL | CHECK | CHECK |
| Dorsomorphin | 1μM | Tocris | 3093 |

**Table S3: E6+10% FBS components.**

| E6+10%FBS | Concentration | Company | Catalogue# |
|---|---|---|---|
| Essential 6 | | Gibco | A1516401 |
| FBS | 10% | Moregate | FBSF |
| Pen-Strep-Neomycin | 1% | Gibco | 15640055 |

**Table S4: List of primers used in this study.**

| Probe gene target | Assay ID | Company | Catalogue# | Notes |
|---|---|---|---|---|
| *POU5F1* | Hs00999632_g1 | Thermo. | 4331182 | See figure 2 |
| *SOX2* | Hs01053049_s1 | Thermo | 4331182 | See figure 2 |
| *NANOG* | Hs02387400_g1 | Thermo | 4331182 | See figure 2 |
| *PDGFRB* | Hs01019589_m1 | Thermo | 4331182 | See figure 2 |
| *CSPG4* | Hs00361541_g1 | Thermo | 4331182 | See figure 2 |
| *ANPEP* | Hs00174265_m1 | Thermo | 4331182 | See figure 2 |
| *ACTA2* | Hs00426835_g1 | Thermo | 4331182 | See figure 2 |
| *FOXF2* | Hs00230963_m1 | Thermo | 4331182 | See figure 2 |
| *FOXC1* | Hs00559473_s1 | Thermo | 4331182 | See figure 2 |
| *VTN* | Hs00940758_g1 | Thermo | 4331182 | See figure 2 |
| *NGFR* | Hs00609976_m1 | Thermo | 4331182 | See Figure S2 |
| *PAX3* | Hs00240950_m1 | Thermo | 4331182 | See Figure S3 |
| *PAX7* | Hs00242962_m1 | Thermo | 4331182 | See Figure S3 |
| *ABCC9* | Hs00245832_m1 | Thermo | 4331182 | See Figure S4 |

Thermo. = Thermo Fisher Scientific

**Table S5: List of inflammatory treatments used in this study.**

| Treatment | Max Conc. (M) | Max Conc. (ng/mL) | Vehicle | Supplier | Cat. # |
|---|---|---|---|---|---|
| IL-1β | $2.89 \times 10^{-10}$ | 5 | PBS + 0.1% BSA | Peprotech | 200-01B |
| TNF | $2.89 \times 10^{-10}$ | 5 | PBS + 0.1% BSA | Peprotech | 300-01A |

**Table S6: List of antibody dilutions used in this study.**

| Stain | Dilution | Cat No. | Company | Secondary | Secondary Dilution | Notes |
|---|---|---|---|---|---|---|
| Hoechst | 1:10,000 | H3570 | Invitrogen | n/a | n/a | n/a |
| OCT3/4 | 1:500 | MAB4401 | Merck | 594 Donkey α Mouse | 1:500 | See figure 2 |
| SOX2 | 1:500 | MAB4360 | Merck | 488 Donkey α Rabbit | 1:500 | See figure 2 |
| NANOG | 1:500 | ab109250 | Abcam | 488 Donkey α Rabbit | 1:500 | See figure 2 |
| PDGFRβ | 1:500 | AF1042 | R&D Systems | 594 Donkey α Goat | 1:500 | See figure 2 |
| CD13 | 1:250 | 301702 | Biolegend | 594 Donkey α Mouse | 1:500 | See figure 2 |
| αSMA | 1:500 | A5228 | Sigma-Aldrich | 594 Donkey α Mouse | 1:500 | See figure 2 |
| NFκB | 1:500 | sc-8008 | Santa Cruz Biotechnology | 594 Donkey α Mouse | 1:500 | See figure 3/5 |
| STAT1 | 1:500 | 14994 | Cell Signalling Technology | 488 Donkey α Rabbit | 1:500 | See figure 4/6 |
| p75 | 1:100 | MA5-13314 | Thermofisher Scientific | 594 Donkey α Mouse | 1:500 | See Figure S2 |
| SMAD2/3 | 1:500 | sc-133098 | Santa Cruz Biotechnology | 594 Donkey α Mouse | 1:500 | See Figures S5-S6 |

**Table S7: List of CBA kits used in this study.**

| CBA Kit target | Catalogue# | Bead Position |
|---|---|---|
| MCP-1 | 558287 | D8 |
| Fractalkine | 560265 | C6 |
| RANTES | 558324 | D4 |
| IL-6 | 558276 | A7 |
| IL-8 | 558277 | A9 |
| TNF | 558273 | D9 |
| sICAM-1 | 560269 | A4 |
| sVCAM-1 | 560427 | D6 |
| VEGF | 558336 | B8 |

**Table S8: Macro used to determine nuclear localisation of immunostaining imaged on the EVOS FL Auto.**

```
//This macro was written by Jacqueline Ross, Biomedical Imaging Research Unit, The University of Auckland, with edits from Sam McCullough.
//This macro opens each RGB stack file in a folder, splits channels and discards either the red or green channel.
//The C3 channel (nuclei) is thresholded, a binary mask created, followed by watershed and the particle analyser.
//The total number of nuclei is counted.
//The regions of interest from the nuclei are stored in the ROI Manager.
//ROI are shown on the C1 image (red) and a threshold selected.
//All ROI are measured. Results and Summary files are saved.
//DAPI image is saved with ROIs and ROI numbers. Protein image is saved as binary mask.
//Images are saved with entire name including .tif. No characters are removed.
//Batch is set to true for maximum speed. No intermediate processing can be viewed.
//Thanks to Phillippe CARL for assistance.
dir1 = getDirectory("Choose Source Directory ");
dir2 = getDirectory("Select Destination  Directory ");
list = getFileList(dir1);
setBatchMode(true);
for (i=0; i<list.length; i++) {
showProgress(i+1, list.length);
open(dir1+list[i]);
imgName=getTitle();
run("Split Channels");
selectWindow("C1-" + imgName);
close();
selectWindow("C3-" + imgName);
run("Options...", "iterations=1 count=1 black");
//run("Threshold...");
setAutoThreshold("IsoData dark");
//setThreshold(22, 255);
setOption("BlackBackground", true);
run("Convert to Mask");
run("Watershed");
run("Set Measurements...", "area mean standard modal min integrated median area_fraction limit display redirect=None decimal=3");
//run("Threshold..."). Lower threshold value is set to select staining above background for protein but also works for binary image.
setAutoThreshold("IsoData dark");
run("Analyze Particles...", "size=30-Infinity summarize add");
selectWindow("C2-" + imgName);
setThreshold(45, 255);
//setThreshold(22, 255);
setOption("BlackBackground", true);
run("Convert to Mask");
//run("Threshold..."). Lower threshold value is set to select staining above background for protein but also works for binary image.
setThreshold(45, 255);
run("Analyze Particles...", "size=1-Infinity summarize add");
roiManager("delete");
//DAPI image is saved with ROIs and labels.
selectWindow("C3-" + imgName);
title = getTitle();
```

```
newtitle= imgName +"-DAPI";
run("Rename...", "title=[" +newtitle+"]");
saveAs("TIFF", dir2 +newtitle);
close();
//Protein image is saved as a binary mask, showing where the labelling was located.
selectWindow("C2-" + imgName);
title = getTitle();
newtitle= imgName +"-GREEN";
run("Rename...", "title=[" +newtitle+"]");
saveAs("TIFF", dir2 +newtitle);
close();
//The results table and the summary table are saved into the Destination directory.
}
{
selectWindow("Summary");
saveAs("Results", dir2 + "Summary.xls");
run("Close");
}
```

**Table S9: Summary of results from each concentration-response experiment.**

| | Day 21 iPSC-derived pericytes | | Day 42 iPSC-derived pericytes | | HPF Pericytes | |
|---|---|---|---|---|---|---|
| | **EC50 (pM)** | **% Responsive cells** | **EC50 (pM)** | **% Responsive cells** | **EC50 (pM)** | **% Responsive cells** |
| | **NFκB** | | **NFκB** | | **NFκB** | |
| **IL-1β** | 7.57 ± 1.14 | 56.8% ± 27.0% | 8.31 ± 3.98 | 58.1% ± 12.9% | 1.75 ± 1.37 | 82.6% ± 12.4% |
| **Exp.1** | 6.76 | 75.95% | 5.49 | 67.15% | 0.587 | 79.46% |
| **Exp.2** | 8.37 | 37.72% | 11.12 | 48.96% | 3.47 | 70.49% |
| **Exp.3** | | | | | 0.702 | 80.54% |
| **Exp.4** | | | | | 2.26 | 100% |
| **TNF** | 7.73 ± 9.09 | 46.1% ± 7.63 | 8.81 | 49.0% | 4.90 ± 3.18 | 77.2% ± 6.83% |
| **Exp.1** | 14.16 | 51.51% | 8.81 | 49.0% | 8.18 | 69.76% |
| **Exp.2** | 1.3 | 40.72% | | | 4.69 | 83.18% |
| **Exp.3** | | | | | 1.84 | 78.68% |
| | **STAT1** | | **STAT1** | | **STAT1** | |
| **IL-1β** | 6.14 ± 7.63 | 50.2% ± 26.2% | n/a | | 3.57 ± 0.87 | 26.9% ± 2.58 |
| **Exp.1** | 0.7407 | 68.74% | n/a | n/a | 4.19 | 24.21% |
| **Exp.2** | 11.53 | 31.62% | n/a | n/a | 2.573 | 29.36% |
| **Exp.3** | | | | | 3.945 | 27.22% |
| **TNF** | n/a | | n/a | | 3.57 ± 0.87 | 18.3% ± 2.05% |
| **Exp.1** | n/a | n/a | n/a | n/a | 0.289 | 20.12% |
| **Exp.2** | n/a | n/a | | | 1.873 | 16.07% |
| **Exp.3** | | | | | 0.6105 | 18.70% |

n/a indicates an experiment which did not show a response. Averaged data is shown ± standard deviation.

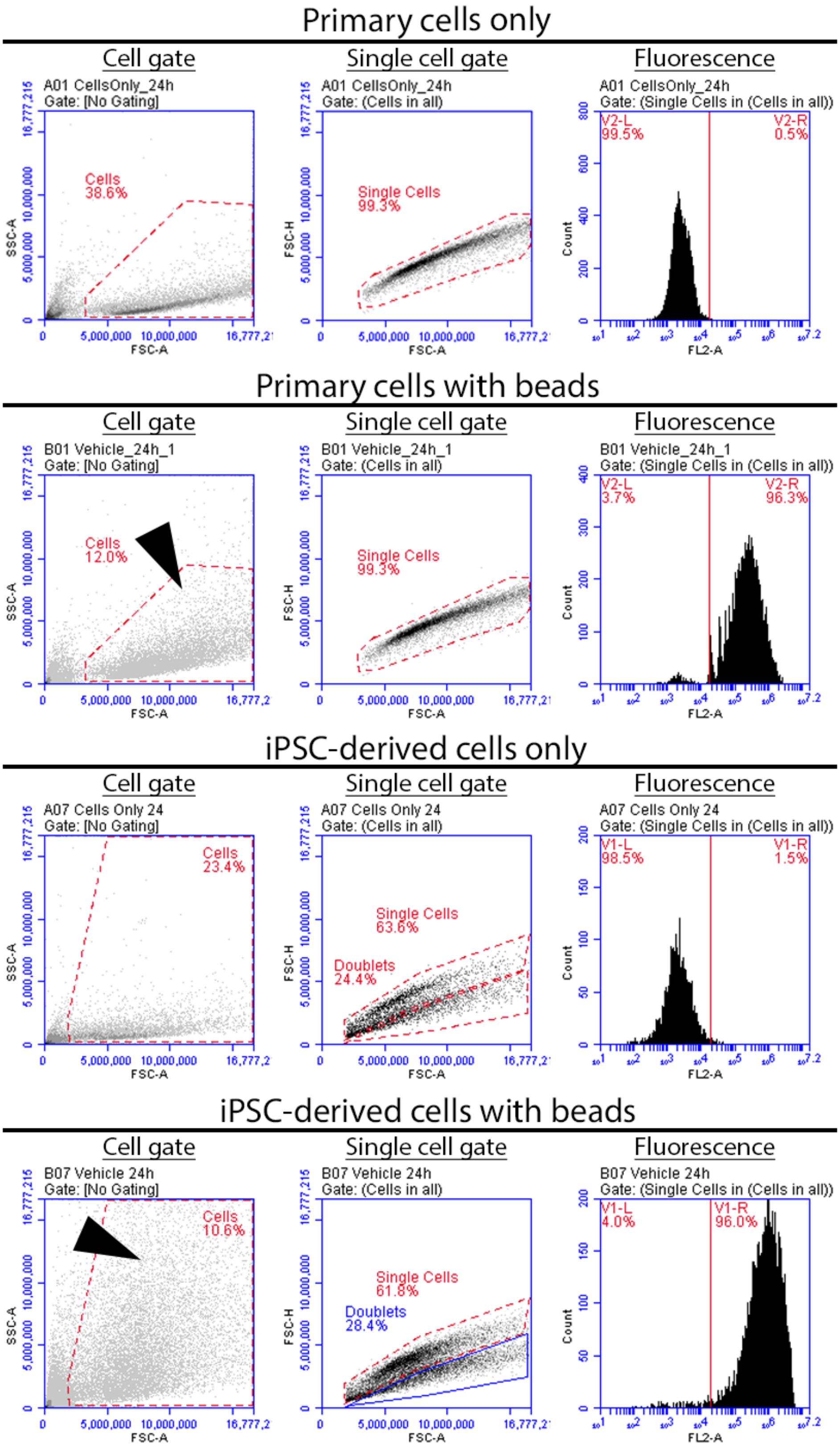


**Figure S1: Gating strategy for primary and iPSC-derived pericytes with and without phagocytosis beads.** Cells are identified by forward (FSC-A) and side scatter (SSC-A) profile. A large inclusion gate is required to collect the more complex form of cells which have phagocytosed beads (arrows). Single cells are identified by comparing forward scatter with height (FSC-H). The threshold for auto-fluorescence is denoted by the vertical red line.

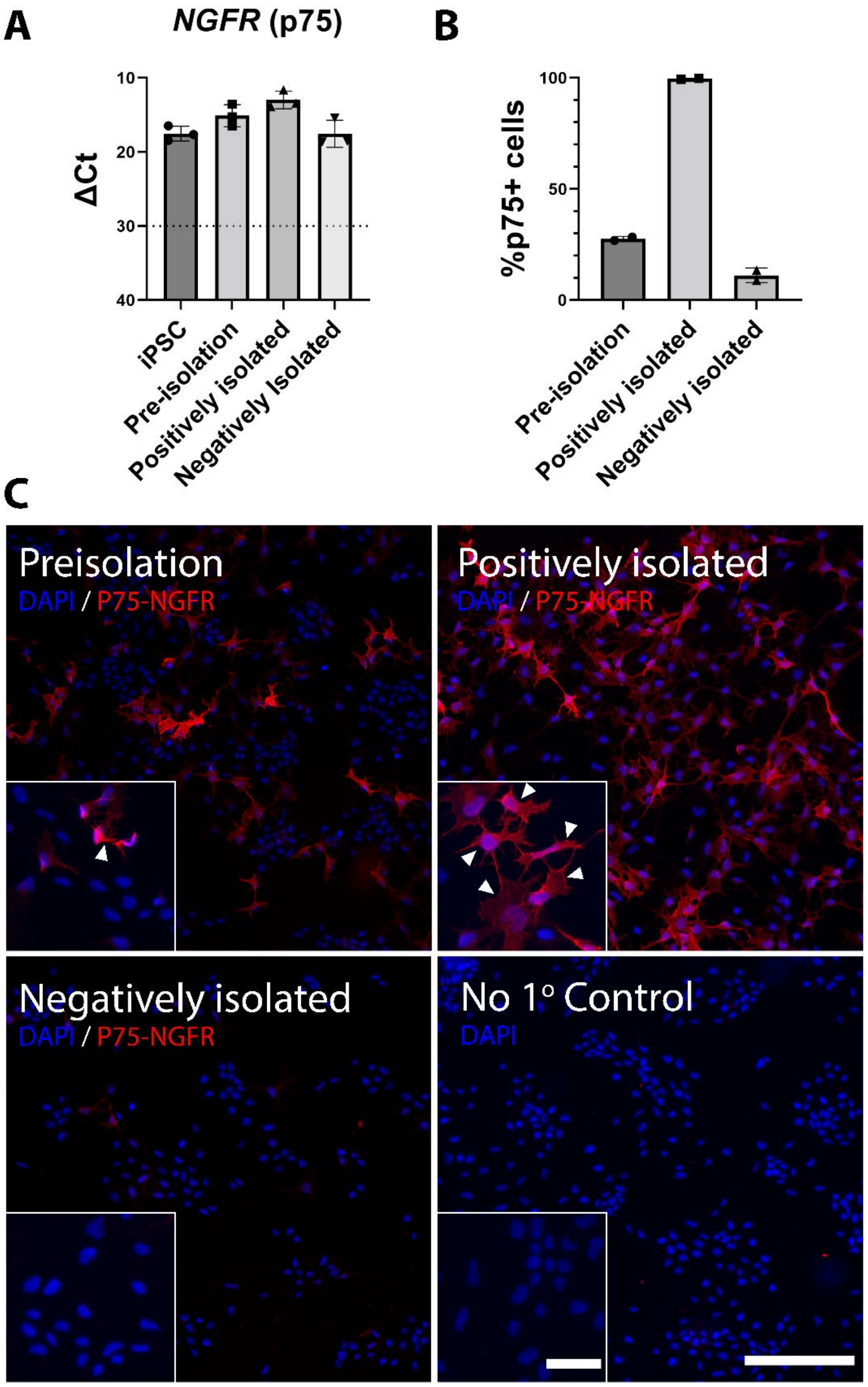


**Figure S2: p75-MACS results in a 99% pure population of neural crest stem cells**. (A) RT-qPCR shows high levels of *NGFR* (p75-NGFR) gene expression in iPSCs and neural crest stem cell (NCSC, day 15 of differentiation) populations before and after p75-MACS isolation. (B) quantification of immunocytochemical staining shows p75 protein expression is only present in 25% of NCSCs before MACS isolation (pre-isolation) but is increased to 99% after MACS isolation (positively isolated). (C) Representative immunocytochemical images of p75 staining in NCSC cell populations. The dotted line on RT-qPCR graphs indicates a ΔCt of 30, demonstrating the minimum ΔCt threshold of expression in these experiments. "No 1° Control" refers to the negative control that didn't receive primary antibody. Scale bar = 200μm in non-magnified images and 50μm in magnified images. Error bars represent standard deviation between the 2-3 experimental repeats.

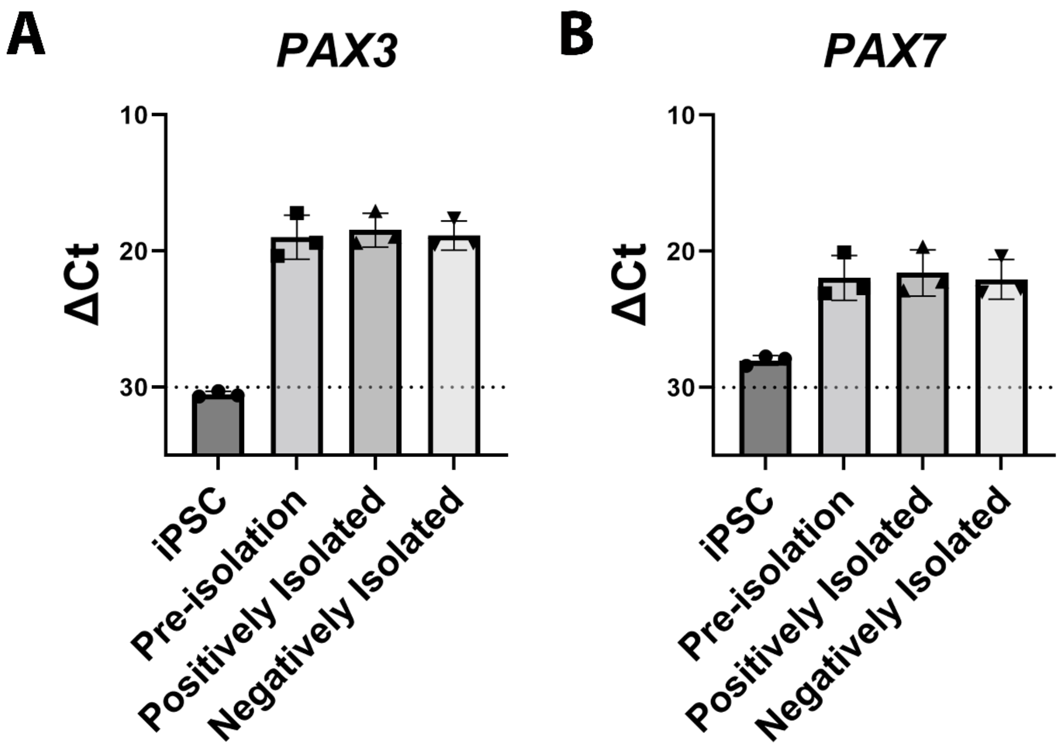


**Figure S3: Neural crest stem cells exhibit expression of *PAX3* and *PAX7*.** (A) RT-qPCR shows high levels of expression of neural crest genes (*PAX3* and *PAX7*) in neural crest stem cell (NCSC, day 15 of differentiation) populations before and after p75-MACS isolation. No PAX3 expression and very low levels of PAX7 expression is observed in iPSCs. The dotted line on RT-qPCR graphs indicates a ΔCt of 30, demonstrating the minimum ΔCt threshold of expression in these experiments. Error bars represent standard deviation between the three experimental repeats.

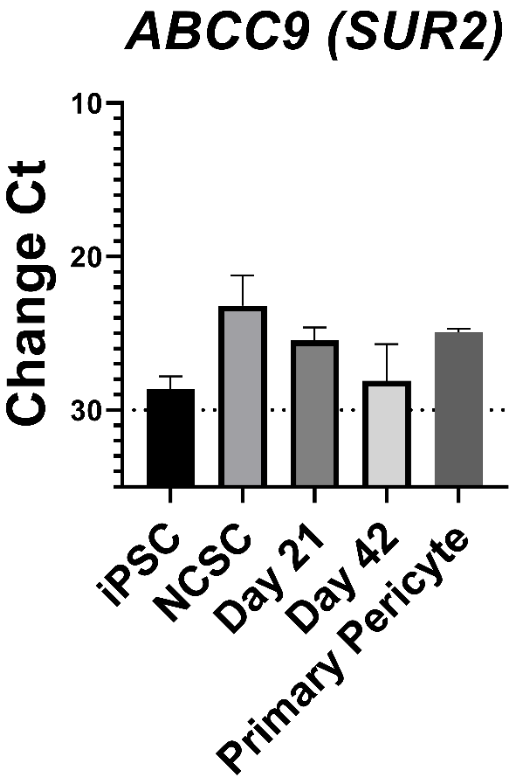


**Figure S4: iPSC-derived brain pericytes exhibit expression of *ABCC9.***

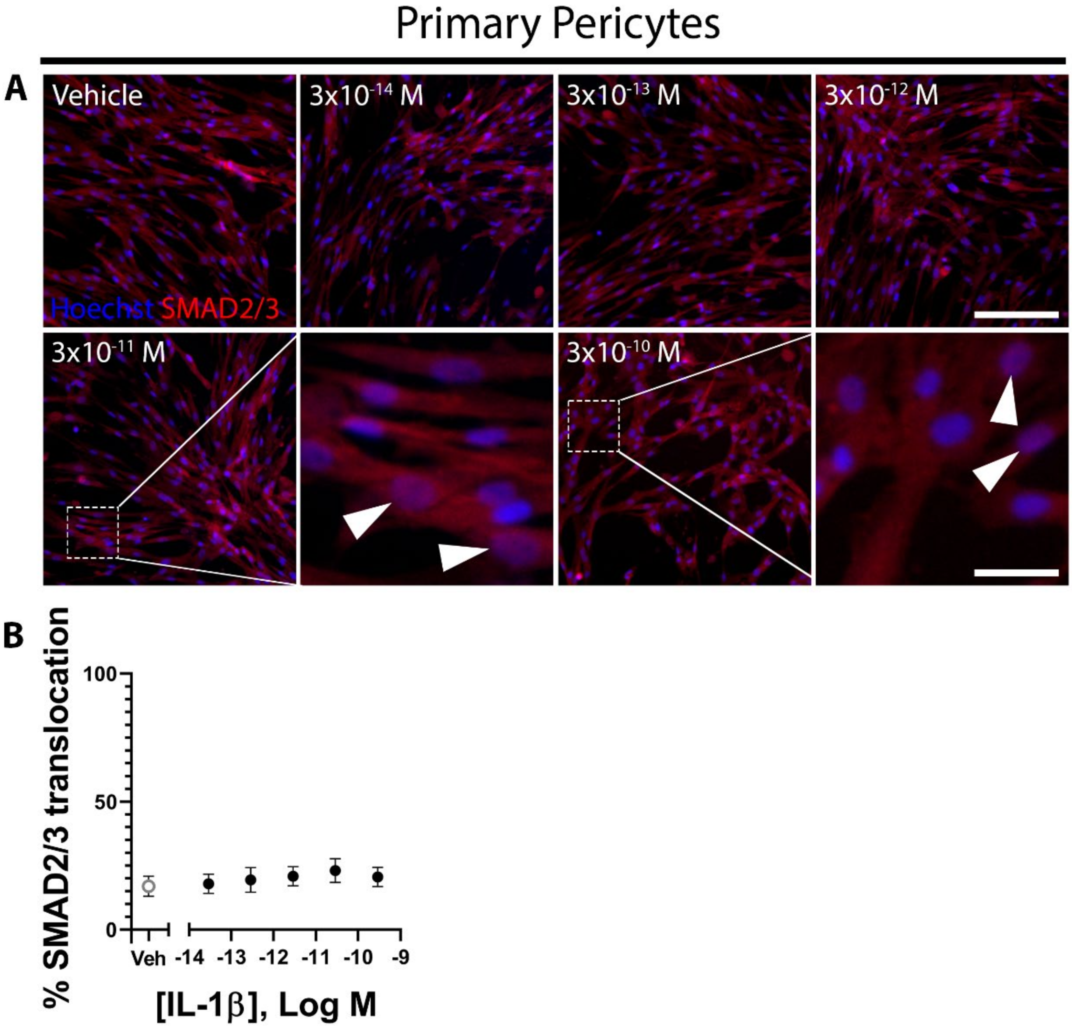


**Figure S5: SMAD2/3 does not translocate to the nucleus with IL-1β treatment.**

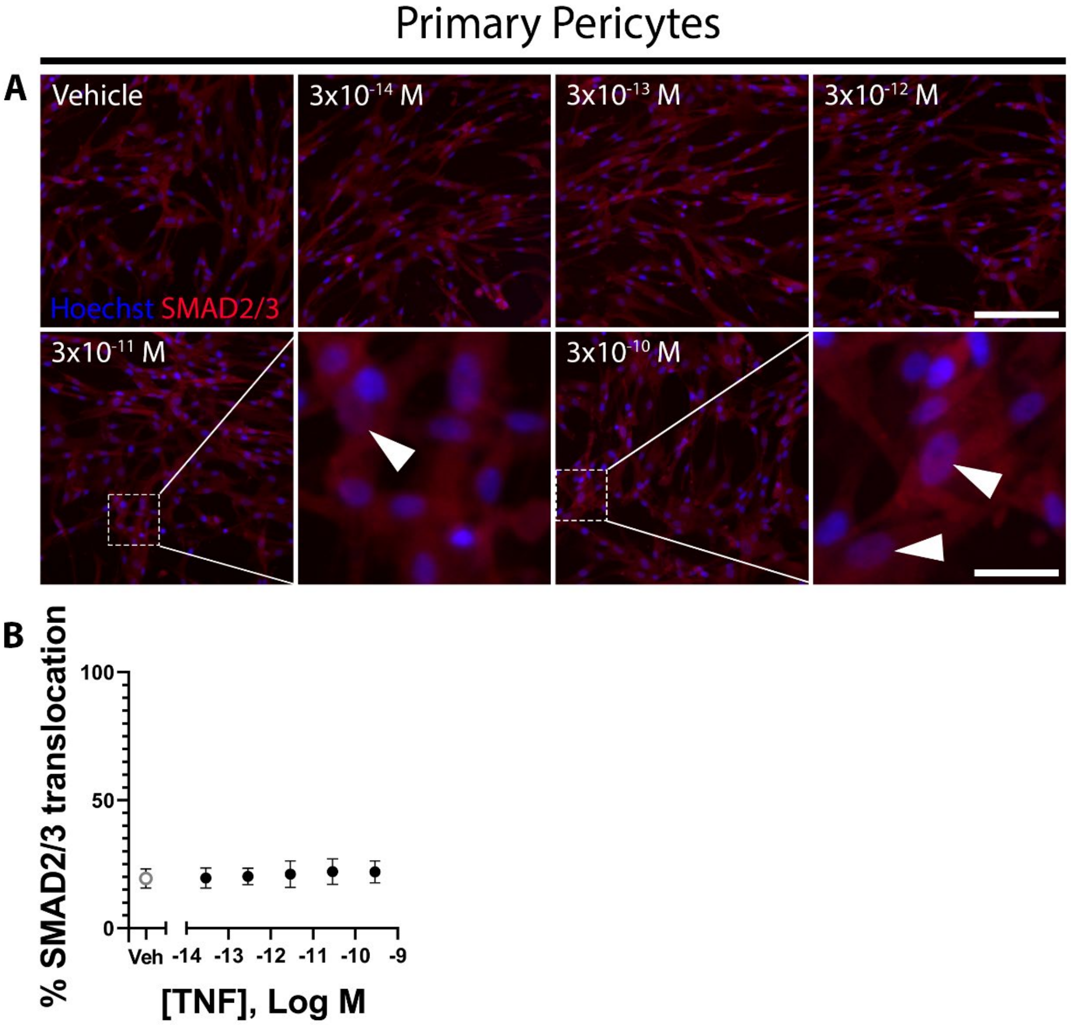


**Figure S6: SMAD2/3 does not translocate to the nucleus with TNF treatment.**

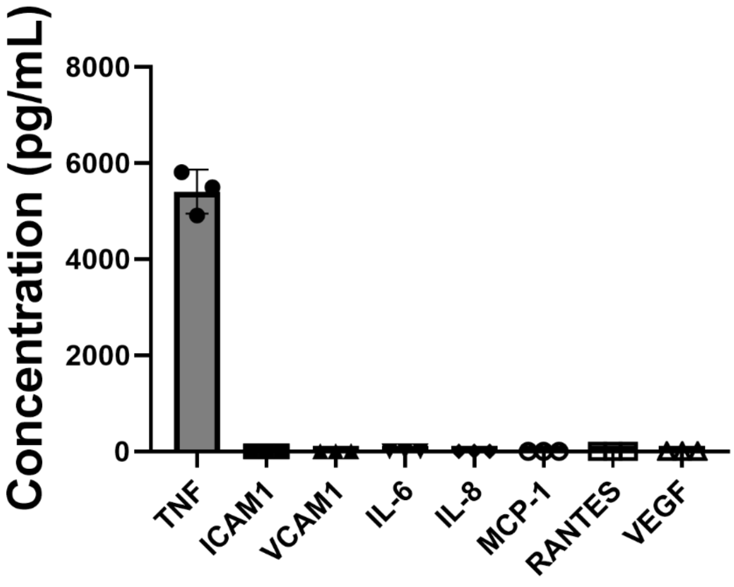


**Figure S7: TNF does not degrade in media only after 72 hours.** Graphs demonstrating the concentration of TNF, sICAM-1, sVCAM, IL-6, IL-8, MCP-1, RANTES, and VEGF in media containing 5000pg/mL TNF after 72 hours. Data presented is averaged from three experimental repeats.
